# Repeated administration of 2-hydroxypropyl-β-cyclodextrin (HPβCD) attenuates the chronic inflammatory response to experimental stroke

**DOI:** 10.1101/2021.05.03.442388

**Authors:** Danielle A. Becktel, Jacob C. Zbesko, Jennifer B. Frye, Amanda G. Chung, Megan Hayes, Kylie Calderon, Jeffrey W. Grover, Anna Li, Frankie G. Garcia, Marco A. Tavera-Garcia, Rick G. Schnellmann, Hsin-Jung Joyce Wu, Thuy-Vi V. Nguyen, Kristian P. Doyle

## Abstract

Globally, more than 67 million people are living with the effects of ischemic stroke. Importantly, many stroke survivors develop a chronic inflammatory response that may contribute to cognitive impairment, a common and debilitating sequela of stroke that is insufficiently studied and currently untreatable. 2-hydroxypropyl-β-cyclodextrin (HPβCD) is an FDA-approved cyclic oligosaccharide that can solubilize and entrap lipophilic substances. The goal of the present study was to determine whether the repeated administration of HPβCD curtails the chronic inflammatory response to stroke by reducing lipid accumulation within stroke infarcts in a distal middle cerebral artery occlusion mouse model of stroke. To achieve this goal, we subcutaneously injected young adult and aged male mice with vehicle or HPβCD three times per week, with treatment beginning one week after stroke. We evaluated mice at 7 weeks following stroke using immunostaining, RNA sequencing, lipidomics, and behavioral analyses. Chronic stroke infarct and peri-infarct regions of HPβCD-treated mice were characterized by an upregulation of genes involved in lipid metabolism and a downregulation of genes involved in innate and adaptive immunity, reactive astrogliosis, and chemotaxis. Correspondingly, HPβCD reduced the accumulation of lipid droplets, T lymphocytes, B lymphocytes, and plasma cells in stroke infarcts. Repeated administration of HPβCD also preserved NeuN immunoreactivity in the striatum and thalamus and c-Fos immunoreactivity in hippocampal regions. Additionally, HPβCD improved recovery through the protection of hippocampal-dependent spatial working memory and reduction of impulsivity. These results indicate that systemic HPβCD treatment following stroke attenuates chronic inflammation and secondary neurodegeneration and prevents post-stroke cognitive decline.

**Significance Statement:** Dementia is a common and debilitating sequela of stroke. Currently, there are no available treatments for post-stroke dementia. Our study shows that lipid metabolism is disrupted in chronic stroke infarcts, which causes an accumulation of uncleared lipid debris and correlates with a chronic inflammatory response. To our knowledge, these substantial changes in lipid homeostasis have not been previously recognized or investigated in the context of ischemic stroke. We also provide a proof of principle that solubilizing and entrapping lipophilic substances using HPβCD could be an effective strategy for treating chronic inflammation after stroke and other CNS injuries. We propose that using HPβCD for the prevention of post-stroke dementia could improve recovery and increase long-term quality of life in stroke sufferers.

## Introduction

Ischemic stroke is a leading cause of death and disability worldwide (Virani et al., 2020). Of stroke survivors, approximately one-third develop a delayed and progressive form of cognitive decline (Black, 2011); however, there are currently no targeted pharmacological interventions to prevent chronic neurodegeneration after stroke. With an aging global population, there is an increasing need for novel and effective treatments that promote recovery after stroke.

We previously reported, in a mouse model of post-stroke dementia, that delayed cognitive impairment is caused by a chronic inflammatory response to stroke that is mediated in part by B lymphocytes (Doyle et al., 2015, 2012). In addition to the progressive infiltration of B lymphocytes, T lymphocytes, and IgA+ plasma cells into the infarct in the weeks following stroke, the chronic inflammatory response is also characterized by the production of neurotoxic molecules such as antibodies, cytokines, and degradative enzymes (Zbesko et al., 2021, 2018). These neurotoxic molecules permeate the glial scar and promote chronic inflammation and secondary neurodegeneration in the surrounding parenchyma (Zbesko et al., 2018). Notably, the chronic inflammatory response to stroke has a similar cellular and molecular profile to atherosclerosis. Chronic stroke infarcts contain foamy macrophages, lipid droplets, and intracellular and extracellular cholesterol crystals (Chung et al., 2018). In atherosclerosis, these distinguishing characteristics are caused by overwhelmed lipid processing systems within macrophages, resulting in the recruitment of adaptive immune cells and the production of pro-inflammatory cytokines and degradative enzymes.

Lipids are principal structural components of the myelin sheath and are major constituents of the human brain (Vance, 2012). Therefore, it is likely that foamy macrophages and cholesterol crystals form in infarcts following ischemic stroke because lipid debris derived from the breakdown of myelin and other cell membranes overwhelms the processing capacity of infiltrating macrophages and resident microglia (Cantuti-Castelvetri et al., 2018). 2-hydroxypropyl-β-cyclodextrin (HPβCD) is a U.S. Food and Drug Administration (FDA)-approved compound that entraps lipids and promotes liver X receptor-mediated transcriptional reprogramming in macrophages to improve cholesterol efflux and incite anti-inflammatory mechanisms (Zimmer et al., 2016). Importantly, HPβCD reduces lipid levels in the lesion following spinal cord injury (Mar et al., 2016) and prevents lipid overload within phagocytic cells in atherosclerosis and Niemann–Pick disease type C (Taylor et al., 2012; Matsuo et al., 2013; Zimmer et al., 2016). Thus, administration of HPβCD is a potential multi-pronged approach for mitigating lipid accumulation in phagocytic cells following ischemic stroke.

Therefore, the objective of this study was to investigate the efficacy of HPβCD as a prospective treatment for chronic inflammation and delayed cognitive impairment following ischemic stroke. To accomplish this objective, the aims of the study were 2-fold: (1) to characterize the lipidome of chronic stroke infarcts and (2) to determine whether lipid complexation and macrophage reprogramming within infarcts, via the repeated systemic administration of HPβCD, attenuates chronic inflammation and secondary neurodegeneration following stroke. To accomplish these aims, we used the distal middle cerebral artery occlusion + hypoxia mouse model of ischemic stroke in conjunction with immunohistochemistry, RNA-Seq, lipidomics, and behavioral analyses. Our resulting data indicate that, coincident with the substantial accumulation of infiltrating immune cells (Doyle et al., 2015; Zbesko et al., 2021), chronic stroke infarcts amass lipids, including sphingomyelins, cholesterol esters, and sulfatides. We also demonstrate that repeated administration of HPβCD in young adult and aged mice attenuates immune cell and lipid accumulation within chronic stroke infarcts and improves recovery at transcriptional and functional levels. Therefore, HPβCD has the potential to improve stroke recovery and prevent post-stroke dementia in humans.

## Materials and Methods

### Experimental design and statistical analysis

The study was designed in a manner consistent with STAIR and RIGOR guidelines (Landis et al., 2012; Lapchak et al., 2012; STAIR, 1999). Sample sizes were determined using power analyses based on expected variances and group differences. Allocation of mice into treatment groups was performed randomly, and behavior experiments were performed with the experimenter blind to treatment group. Statistical analyses were performed with Prism 6.0 (GraphPad). Data are presented as the mean ± standard error of mean (SEM). Group sizes, statistical tests, and *p* values for each experiment are reported in **Table 1**. For CBC and lipidomics analyses, normality was assessed using the Kolmogorov—Smirnov test. For all other analyses, data from each individual mouse is presented so the distribution within each group can be visually evaluated; however, the normality was not formally assessed. For GSEA, the normalized enrichment score (NES) is reported. The NES accounts for differences in gene set size and in correlations between gene sets. The NES is based on all dataset permutations to correct for multiple hypothesis testing.

**Table 1.** Statistical table.

| Figure | Group Size<br>(n) | Statistical<br>Test | Statistical Values | p value | post hoc<br>Comparisons | p value |
| --- | --- | --- | --- | --- | --- | --- |
| 1B | 4 | Paired *t* test | *M* (Contra Cortex -<br>Infarct) = -51.71<br>*SEM* = 1.300 | < 0.0001 | N/A | N/A |
| 1C | Infarct = 7<br>Contra<br>Cortex = 4 | Unpaired *t* test | *M* (Contra Cortex -<br>Infarct) = -34.14<br>*SEM* = 1.077 | < 0.0001 | N/A | N/A |
| 1D | 8 | Multiple *t* tests;<br>Holm-Sidak's<br>multiple<br>comparisons<br>test | N/A | N/A | SM 14:0_16.51<br>SM 15:0_17.67<br>SM 16:0_19.05<br>SM 16:1_16.92<br>SM 16:1_17.23<br>SM 18:0_22.55<br>SM 18:1_19.71<br>SM 18:1_20.12<br>SM 19:0_24.48<br>SM 20:0_26.16<br>SM 20:1_23.41<br>SM 21:0_27.48<br>SM 22:0_28.28<br>SM 22:1_26.30<br>SM 22:1_26.87<br>SM 22:2_23.57<br>SM 23:0_28.82<br>SM 23:1_27.48<br>SM 23:2_25.37<br>SM 24:0_29.21<br>SM 24:1_28.23<br>SM 24:2_26.78<br>SM 24:3_24.26<br>SM 24:3_24.64<br>SM 24:4_22.21<br>SM 25:0_29.62<br>SM 26:0_29.96<br>SM 26:1_29.14<br>SM 26:2_28.42 | 0.000003<br>< 0.000001<br>< 0.000001<br>0.000003<br>0.000004<br>0.00379<br>0.000172<br>0.009913<br>0.002555<br>0.000813<br>0.0018<br>0.006607<br>< 0.000001<br>0.000006<br>0.000056<br>0.000003<br>0.002919<br>< 0.000001<br>0.000017<br>< 0.000001<br>< 0.000001<br>0.000062<br>0.000002<br>0.000004<br>0.000005<br>0.000226<br>0.000076<br>0.000001<br>0.00016 |
| 2A | 4 | Paired *t* test | *M* (Contra Cortex -<br>Infarct) = -54.83<br>*SEM* = 1.527 | < 0.0001 | N/A | N/A |
| 2B | 9 | Paired *t* test | *M* (Contra Cortex -<br>Infarct) = -27.39 | < 0.0001 | N/A | N/A |
|  |  |  | SEM = 0.9784 |  |  |  |
| 2C | 5 | Multiple *t* tests;<br>Holm-Sidak's<br>multiple<br>comparisons<br>test | N/A | N/A | CE 12:0 | 0.007817 |
|  |  |  |  |  | CE 14:0 | 0.000752 |
|  |  |  |  |  | CE 15:0 | 0.003201 |
|  |  |  |  |  | CE 16:0 | 0.000217 |
|  |  |  |  |  | CE 16:1 | 0.000495 |
|  |  |  |  |  | CE 17:0 | 0.001331 |
|  |  |  |  |  | CE 18:0 | 0.000894 |
|  |  |  |  |  | CE 18:1 | 0.001131 |
|  |  |  |  |  | CE 18:2 | 0.001921 |
|  |  |  |  |  | CE 18:3 | 0.001145 |
|  |  |  |  |  | CE 20:0 | 0.005071 |
|  |  |  |  |  | CE 20:3 | 0.001136 |
|  |  |  |  |  | CE 20:4 | 0.000936 |
|  |  |  |  |  | CE 20:5 | 0.000785 |
|  |  |  |  |  | CE 22:0 | 0.00243 |
|  |  |  |  |  | CE 22:6 | 0.005786 |
|  |  |  |  |  | CE 24:0 | 0.007793 |
|  |  |  |  |  | CE 24:1 | 0.000802 |
| 5B (B220) | Vehicle = 11 | Unpaired *t* test | *M* (HPβCD - Vehicle)<br>= -12.22 | < 0.0001 | N/A | N/A |
|  | HPβCD = 10 |  | SEM = 1.819 |  |  |  |
| 5B (CD3ε) | Vehicle = 11 | Unpaired *t* test | *M* (HPβCD - Vehicle)<br>= -7.695 | < 0.0001 | N/A | N/A |
|  | HPβCD = 10 |  | SEM = 0.9035 |  |  |  |
| 5B (CD138) | 4 | Unpaired *t* test | *M* (HPβCD - Vehicle)<br>= -51.22 | 0.0163 | N/A | N/A |
|  |  |  | SEM = 15.49 |  |  |  |
| 5B (IgA) | 8 | Unpaired *t* test | *M* (HPβCD - Vehicle)<br>= -34.30 | 0.0026 | N/A | N/A |
|  |  |  | SEM = 9.362 |  |  |  |
| 5B (ORO) | 4 | Unpaired *t* test | *M* (HPβCD - Vehicle)<br>= -4.999 | 0.0462 | N/A | N/A |
|  |  |  | SEM = 1.995 |  |  |  |
| 5B (Alcian Blue) | 3 | Unpaired *t* test | *M* (HPβCD - Vehicle)<br>= -1.576 | 0.7977 | N/A | N/A |
|  |  |  | SEM = 5.751 |  |  |  |
| 6B (B220) | Vehicle = 4 | Unpaired *t* test | *M* (HPβCD - Vehicle)<br>= -6.580 | 0.0039 | N/A | N/A |
|  | HPβCD = 5 |  | SEM = 0.8794 |  |  |  |
| 6B (CD3ε) | Vehicle = 8 | Unpaired *t* test | *M* (HPβCD - Vehicle)<br>= -12.75 | 0.0002 | N/A | N/A |
|  | HPβCD = 7 |  | SEM = 2.429 |  |  |  |
| 6B (IgA) | Vehicle = 5 | Unpaired *t* test | *M* (HPβCD - Vehicle)<br>= -21.55 | 0.0477 | N/A | N/A |
| | HPβCD = 6 | | SEM = 8.139 | | | |
| 6B (ORO) | 4 | Unpaired *t* test | *M* (HPβCD - Vehicle)<br>= -9.330<br>SEM = 1.124 | 0.0002 | N/A | N/A |
| 6B (Alcian Blue) | Vehicle = 9<br>HPβCD = 10 | Unpaired *t* test | *M* (HPβCD - Vehicle)<br>= -1.372<br>SEM = 1.954 | 0.4921 | N/A | N/A |
| 7B (CD19+GL 7-) | Vehicle = 10<br>HPβCD = 9 | Unpaired *t* test | *M* (HPβCD - Vehicle)<br>= 3.258<br>SEM = 2.553 | 0.2191 | N/A | N/A |
| 7B (CD19+GL 7+) | Vehicle = 10<br>HPβCD = 9 | Unpaired *t* test | *M* (HPβCD - Vehicle)<br>= -0.1013<br>SEM = 0.1402 | 0.4796 | N/A | N/A |
| 7C (IgG1) | Vehicle = 10<br>HPβCD = 9 | Unpaired *t* test | *M* (HPβCD - Vehicle)<br>= -2.589<br>SEM = 28.71 | 0.9292 | N/A | N/A |
| 7C (IgA) | Vehicle = 10<br>HPβCD = 9 | Unpaired *t* test | *M* (HPβCD - Vehicle)<br>= 7.567<br>SEM = 16.23 | 0.647 | N/A | N/A |
| 7E (CD4+ T cells) | 9 | Unpaired *t* test | *M* (HPβCD - Vehicle)<br>= 0.03444<br>SEM = 0.9497 | 0.9715 | N/A | N/A |
| 7E (CD8+ T cells) | 9 | Unpaired *t* test | *M* (HPβCD - Vehicle)<br>= -0.08889<br>SEM = 1.201 | 0.9419 | N/A | N/A |
| 7E (Tregs) | 9 | Unpaired *t* test | *M* (HPβCD - Vehicle)<br>= -1.844<br>SEM = 2.238 | 0.422 | N/A | N/A |
| 7F | 6 | Multiple *t* tests;<br>Holm-Sidak's<br>multiple<br>comparisons<br>test | N/A | N/A | WBC<br>Neutrophils<br>Lymphocytes<br>Monocytes<br>Eosinophils<br>Basophils | 0.109564<br>0.213092<br>0.0445<br>0.027171<br>0.662546<br>0.58393 |
| 8E (*Npc1*) | 4 | Unpaired *t* test | *M* (HPβCD - Vehicle)<br>= 0.8853<br>SEM = 0.1417 | 0.0008 | N/A | N/A |
| 8E (*Npc2*) | 4 | Unpaired *t* test | *M* (HPβCD - Vehicle)<br>= 0.6104<br>SEM = 0.1525 | 0.0071 | N/A | N/A |
| 8E (*Abca1*) | 4 | Unpaired *t* test | *M* (HPβCD - Vehicle)<br>= 0.7311<br>SEM = 0.2348 | 0.0208 | N/A | N/A |
| 8E (*Apoe*) | 4 | Unpaired *t* test | *M* (HPβCD - Vehicle)<br>= 0.6647<br>SEM = 0.2186 | 0.0228 | N/A | N/A |
| 8E (*Trem2*) | 4 | Unpaired *t* test | *M* (HPβCD - Vehicle)<br>= 0.7428 | 0.0187 | N/A | N/A |
|  |  |  | SEM = 0.2325 |  |  |  |
| 14A (Time) | Vehicle = 15 | Unpaired *t* test | *M* (HPβCD - Vehicle) = -26.63 | 0.0393 | N/A | N/A |
|  | HPβCD = 13 |  | SEM = 12.27 |  |  |  |
| 14A (Latency) | Vehicle = 15 | Unpaired *t* test | *M* (HPβCD - Vehicle) = 157.5 | 0.0165 | N/A | N/A |
|  | HPβCD = 13 |  | SEM = 61.44 |  |  |  |
| 14B (% Correct Alternations) | Vehicle = 15 | Unpaired *t* test | *M* (HPβCD - Vehicle) = 9.389 | 0.0060 | N/A | N/A |
|  | HPβCD = 13 |  | SEM = 3.137 |  |  |  |
| 14B (Total Entries) | Vehicle = 15 | Unpaired *t* test | *M* (HPβCD - Vehicle) = -3.385 | 0.1847 | N/A | N/A |
|  | HPβCD = 13 |  | SEM = 2.484 |  |  |  |
| 14C | Vehicle = 7 | Unpaired *t* test | *M* (HPβCD - Vehicle) = -0.1564 | 0.0311 | N/A | N/A |
|  | HPβCD = 9 |  | SEM = 0.06527 |  |  |  |
| 14D | 12 | Pearson correlation analysis | Pearson *r* = -0.5640 | 0.0561 | N/A | N/A |
|  |  |  | R<sup>2</sup> = 0.3181 |  |  |  |
| 14E | 5 | Unpaired *t* test | *M* (HPβCD - Vehicle) = 0.3339 | 0.0487 | N/A | N/A |
|  |  |  | SEM = 0.1438 |  |  |  |
| 14F | 5 | Unpaired *t* test | *M* (HPβCD - Vehicle) = 0.3901 | 0.0039 | N/A | N/A |
|  |  |  | SEM = 0.09717 |  |  |  |
| 14G (CA1 Hippocampus) | Vehicle = 9 | Unpaired *t* test | *M* (HPβCD - Vehicle) = 0.3379 | 0.0268 | N/A | N/A |
|  | HPβCD = 10 |  | SEM = 0.1394 |  |  |  |
| 14G (CA3 Hippocampus) | Vehicle = 9 | Unpaired *t* test | *M* (HPβCD - Vehicle) = 0.3196 | 0.0229 | N/A | N/A |
|  | HPβCD = 10 |  | SEM = 0.1278 |  |  |  |
| 14G (Dentate Gyrus) | Vehicle = 9 | Unpaired *t* test | *M* (HPβCD - Vehicle) = 0.6193 | 0.0178 | N/A | N/A |
|  | HPβCD = 10 |  | SEM = 0.2362 |  |  |  |

### Mice

Young adult (3- to 4-month-old) and aged (17- to 18-month-old) wild-type male C57BL/6J mice (Stock No. 000664) were purchased from The Jackson Laboratory. Mice were housed in a temperature-controlled suite under a 12-hour light–dark regimen, with food and water available *ad libitum*. All procedures met NIH guidelines and were approved by the University of Arizona Institutional Animal Care and Use Committee. At each time point, mice were euthanized by isoflurane anesthesia (JD Medical), exsanguination, and subsequent intracardial perfusion with 0.9% saline. Whole brains or individual brain regions were then removed and either placed in RNAlater (Invitrogen, Cat. No. AM7020) for RNA sequencing analysis; flash frozen in liquid nitrogen for lipidomics analysis; or placed in a 4% paraformaldehyde (PFA) solution for 24 h before being transferred into a 30% sucrose solution for immunostaining analysis. Whole blood collected at the time of euthanasia was either placed directly in heparin-coated tubes for complete blood count analysis or extracted with sodium citrate to separate plasma via centrifugation.

### Stroke surgeries

Stroke was induced in mice using the distal middle cerebral artery occlusion + hypoxia (DH) model. The DH stroke model generates a sizable infarct (24% of the ipsilateral hemisphere centered on the somatosensory cortex), has little variability, and has exceptional long-term survivability (Doyle et al., 2012; Nguyen et al., 2016). To induce stroke, we anesthetized animals by isoflurane inhalation and kept them at 37°C throughout the surgical procedure. For all experiments, we injected mice subcutaneously (s.c.) with a single dose of buprenorphine hydrochloride (0.1 mg/kg; Henry Schein) and a single dose of cefazolin antibiotic (25 mg/kg; Sigma-Aldrich) dissolved in sterile saline. Following pre-operative preparation, an incision was made to expose the right temporoparietal skull between the orbit and the ear. Under an operating microscope, a small hole was made with a high-speed microdrill through the outer surface of the semi-translucent skull over the visually identified middle cerebral artery (MCA) at the level of the parietal cerebral artery. Permanent occlusion of the MCA was performed by electrocoagulation with a small vessel cauterizer (Bovie Medical Corporation). Surgical wounds were closed using Surgi-lock 2oc tissue adhesive (Meridian Animal Health). We then immediately transferred mice to a hypoxia chamber (Coy Laboratory Products) containing 9% oxygen and 91% nitrogen for 45 min. Sustained-release buprenorphine (Bup-SR, 1 mg/kg s.c.; ZooPharm) was administered 24 h after surgery as post-operative analgesia.

### Magnetic resonance imaging (MRI)

Infarct volume was assessed by magnetic resonance imaging (MRI) 24 h after stroke using a Bruker Biospec 70/20 7.0T scanner with ParaVision-360.2.0 software and a 4-channel phase array mouse coil (Bruker Biospin GmbH, Germany). Mice were placed in a cradle equipped with a stereotaxic frame, an integrated heating system to maintain body temperature at 37±1°C, and a pressure probe to monitor respiration. During MRI acquisition, anesthesia was maintained by inhalation of 1.5-3% isoflurane.

High-resolution structural images were acquired using a *T_2_*-weighted RARE multi-echo Bruker pulse sequence with the following parameters: repetition time (*TR*) = 2000 ms; flip angle = 180°; RARE factor = 12; matrix size = 128 x 128; averages = 2; field of view (FOV) = 1.92 cm x 1.92 cm; slice thickness = 0.8 mm; number of slices = 15; acquisition time = 8 min 53 s. Infarcts and hemispheric cross sections were manually delineated on *T_2_*-weighted images using Mango v4.1.

### Drug treatment

We injected HPβCD or vehicle s.c. three times per week, beginning 7 days after stroke surgery, for 6 weeks until euthanasia at 7 weeks after stroke. Mice in the treatment group received 2-hydroxypropyl-β-cyclodextrin powder (Sigma-Aldrich, Cat. No. H-107) dissolved in sterile phosphate-buffered saline (PBS) at a dose of 4 g HPβCD/kg body weight. Mice in the vehicle control group received 300 μL sterile PBS.

### Immunostaining

Coronal brain sections (40 μm) were collected using a freezing Microm HM 450 sliding microtome (Thermo Fisher Scientific) and stored in cryoprotectant medium at –20°C until processing. Immunostaining was performed on free-floating brain sections using standard protocols (Doyle et al., 2015; Nguyen et al., 2016; Zbesko et al., 2018). Primary antibodies against neuronal nuclei (NeuN; 1:500; Millipore Sigma, Cat. No. MAB377; RRID:AB_2298772), c-Fos (1:2000; Abcam, Cat. No. ab190289; RRID:AB_2737414), CD3ε (1:1000; BD Biosciences, Cat. No. 550277; RRID:AB_393573), B220/CD45R (1:500; BD Biosciences, Cat. No. 553085; RRID:AB_394615), immunoglobulin A (IgA; 1:1000; BioLegend, Cat. No. 407004; RRID:AB_315079), and CD138 (Syndecan-1; 1:200; BioLegend, Cat. No. 142514; RRID:AB_2562198) were used in conjunction with the appropriate secondary antibody and visualized using the VECTASTAIN Elite ABC Reagent, Peroxidase, R.T.U. (Vector Laboratories, Cat. No. PK-7100) and Vector DAB Substrate (3,3’-diaminobenzidine) Kit (Vector Laboratories, Cat. No. SK-4100). Secondary antibodies were diluted 1:400 for biotinylated horse anti-mouse IgG (Vector Laboratories, Cat. No. BA-2000; RRID:AB_2313581), biotinylated goat anti-hamster IgG (Vector Laboratories, Cat. No. BA-9100; RRID:AB_2336137), and biotinylated rabbit anti-rat IgG (Vector Laboratories, Cat. No. BA-4000; RRID:AB_2336206). Sections were imaged using a Keyence BZ-X700 digital microscope with phase contrast, light, and fluorescence capabilities.

### Oil Red O staining

Frozen tissue sections were mounted on slides and allowed to dry. Mounted sections were dehydrated in 100% propylene glycol for 5 min and then stained with Oil Red O (ORO) for 10 min (Abcam, Cat. No. ab150678). The sections were then differentiated in 85% propylene glycol for 3 min and rinsed with distilled water three times. Coverslips were applied to all slides using an aqueous mounting medium (Vector Laboratories, Cat. No. H-1400). Stained sections were imaged with a Keyence BZ-X700 digital microscope.

### Alcian blue staining

Frozen tissue sections were mounted on slides and allowed to dry. Slides were immersed in eosin (VWR, Cat. No. 95057-848) and then rinsed eight times in PBS. Slides were incubated in 3% acetic acid (pH 2.5) for 3 min and then placed in a solution of 1% w/v Alcian blue in 3% acetic acid (pH 2.5) for 30 min (EMD Millipore, Cat. No. TMS-010-C). Slides were then rinsed in 3% acetic acid and allowed to dry overnight. Slides were cleared in xylene and preserved with Entellan mounting medium (EMD Millipore, Cat. No. 14802) and coverslips. Stained sections were imaged with a Keyence BZ-X700 digital microscope.

### Quantification of staining

To quantify the amount of positive staining for CD3ε, B220, Oil Red O, and Alcian blue, we used Fiji (Schindelin et al., 2012). The ‘Colour Deconvolution’ function was used to separate color channels in images of Alcian blue staining. We then converted all stitched images to 8-bit and traced the stroke infarct using the ‘Polygon selections’ tool. We applied a threshold of positive staining to each image and measured the pixel area of positive staining within the selected region. To quantify IgA and CD138, we utilized the ‘Cell Counter’ plugin to manually count the number of positive cells within the stroke infarct. To quantify NeuN and c-Fos immunoreactivity in vehicle- and HPβCD-treated mice, we used Fiji to perform global thresholding in conjunction with a watershed algorithm on the ipsilateral side and equivalent region in the contralateral hemisphere. Fields covering the selected regions were analyzed at bregma +0.38 mm and −1.46 mm (NeuN) or −1.82 mm (c-Fos) using a 20× objective lens. We also measured hippocampal area in images captured with a 2× objective lens by tracing the hippocampus with the ‘Polygon selections’ tool in Fiji.

### RNA sequencing and data analysis

Infarcts were visually identified and dissected using forceps and a razor blade. Following infarct removal, adjacent peri-infarct regions were dissected. Corresponding regions in the contralateral hemisphere were also dissected. Fresh brain tissue collected from vehicle- and HPβCD-treated mice was immersed in RNAlater (Invitrogen, Cat. No. AM7020) and delivered to the University of Arizona Genetics Core. Samples were assessed for quality with an Advanced Analytics Fragment Analyzer (High Sensitivity RNA Analysis Kit, Cat. No. DNF-491/User Guide DNF-491-2014AUG13) and quantity with a Qubit RNA HS Assay Kit (Cat. No. Q32852). Given satisfactory quality (RNA integrity number >8) and quantity, samples were used for library construction with the TruSeq Stranded mRNA Library Prep Kit from Illumina (Cat. No. 20020595), as well as the KAPA Dual-Indexed Adapter Kit from Roche (Cat. No. 8278555702). Upon completion of library construction, samples were assessed for quality and average fragment size with the Advanced Analytics Fragment Analyzer (High Sensitivity NGS Analysis Kit, Cat. No. DNF-846/User Guide DNF-486-2014MAR10). Quantity was assessed with an Illumina Universal Adaptor-specific qPCR kit from KAPA Biosystems (KAPA Library Quantification Kit for Illumina NGS, Cat. No. KK4824/KAPA Library Quantification Technical Guide – AUG2014). Following the final library quality control, samples were equimolar-pooled and clustered for paired-end sequencing on the Illumina NextSeq500 machine to generate 75 bp reads. The sequencing run was performed using Illumina NextSeq500 run chemistry (NextSeq500/550 High Output v2 Kit 150 cycles, Cat. No. FC-404-2002). Sequencing data is publicly available at the National Center for Biotechnology Information through Gene Expression Omnibus accession numbers: GSE173544 and GSE173715. For data analysis, the resulting sequences were demultiplexed using bcl2fastq v2.19 (Illumina) and trimmed of their indexing adaptors using Trimmomatic v0.32 (Bolger, Lohse, & Usadel, 2014). The trimmed reads were aligned to the GRCm38 reference genome using STAR v2.5.2b (Dobin et al., 2013). Gene expression was calculated using the htseq-count function of the HTSeq python tool (Anders, Pyl, & Huber, 2015). Genes were annotated using the BioMart database. Differential expression analysis of count tables was performed using DESeq2 (Love, Huber, & Anders, 2014). Gene set enrichment analysis (GSEA) was performed on all significant differentially expressed genes (false discovery rate [FDR]-adjusted *p* < 0.05) using a database of Gene Ontology (GO) terms for biological processes. Enrichment maps were constructed from GO terms using the Enrichment Map Cytoscape application (Merico, Isserlin, Stueker, Emili, & Bader, 2010). Minimal editing, such as repositioning of nodes and removal of repetitive gene-sets, was performed to optimize the map layout. Pathway analysis was performed on all differentially expressed genes using Ingenuity Pathway Analysis v01-13 (IPA).

### Targeted lipidomics analysis (LIPID MAPS Lipidomics Core)

Fresh brain tissue dissected from vehicle- and HPβCD-treated mice was flash frozen and delivered to the LIPID MAPS Lipidomics Core at the University of California, San Diego. Upon arrival, each sample underwent lipid extraction and quality control analyses. The comprehensive sphingolipid panel and cholesterol ester panel were independently performed using established protocols (Quehenberger et al., 2010).

### Flow cytometry

To assess peripheral immune cell populations, spleens were extracted after transcardial perfusion with 0.9% saline. Each spleen was crushed in 1 mL ammonium–chloride– potassium RBC lysing buffer and resuspended in 1 mL complete DMEM. For cell staining, fluorophore-conjugated monoclonal antibodies specific for CD4 (RM4-5), CD8α (53-6.7), and TCRβ (H57-597) were obtained from BioLegend. Monoclonal antibodies specific for CD19 (6D5) and GL7 (GL7) were purchased from BD Pharmingen. Antibodies against IgG1 (RMG1-1) and IgA (RMA-1) were obtained from BioLegend. For intranuclear staining, buffers from a Foxp3 Staining Buffer Set (eBioscience) were used to stain antibodies recognizing Foxp3 (FJK-16s, eBioscience). Cells were run on a BD LSR II flow cytometer (BD Biosciences) and analyses were performed with FlowJo v10.7 for Windows (Tree Star).

### Y-maze spontaneous alternation behavior test

Spatial working memory was assessed with the spontaneous alternation behavior (SAB) paradigm. Testing occurred in a Y-shaped maze consisting of two symmetrical arms and one longer arm arranged at 120° angles (symmetrical arms: 7.5 cm wide, 37.0 cm long, 12.5 cm high; longer arm: 7.5 cm wide, 42.0 cm long, 12.5 cm high). Mice were placed at the end of the longest arm and allowed to freely explore the three arms in a 5 min trial, as described previously (Doyle et al., 2015). The number of arm entries and the number of correct alternations were recorded by an ANY-maze behavioral video tracking software (Stoelting, Co.). An entry was recorded when all four limbs of the mouse were within an arm.

### Light/dark transition test

The light/dark box arena consisted of a Plexiglas box (40 cm wide, 40 cm long, 35 cm high) divided by a small underpass into two equally sized compartments: a brightly illuminated zone (390 lux) and a covered dark zone (2 lux). Prior to the test, animals were habituated to the dark for 30 min. Mice were then placed in the dark chamber of the arena and allowed to move freely between the two chambers for 10 min. The time spent in each chamber, the latency to exit the dark chamber, and the total number of transitions were recorded by an ANY-maze behavioral video tracking software (Stoelting, Co.).

## Results

### Lipid composition of chronic stroke infarcts in young adult mice

Prior to evaluating the lipid composition of chronic stroke infarcts, we performed *T_2_*-weighted MRI on young adult mice 24 hours after stroke to visualize the acute area of infarction generated by the distal middle cerebral artery occlusion + hypoxia (DH) stroke model. As previously shown, infarcts were reliably centered on the somatosensory cortex and included a portion of the M1 region of the primary motor cortex (Doyle et al., 2012). Notably, infarcts did not extend to the hippocampus (**Fig. 1A**).

**Figure 1.**
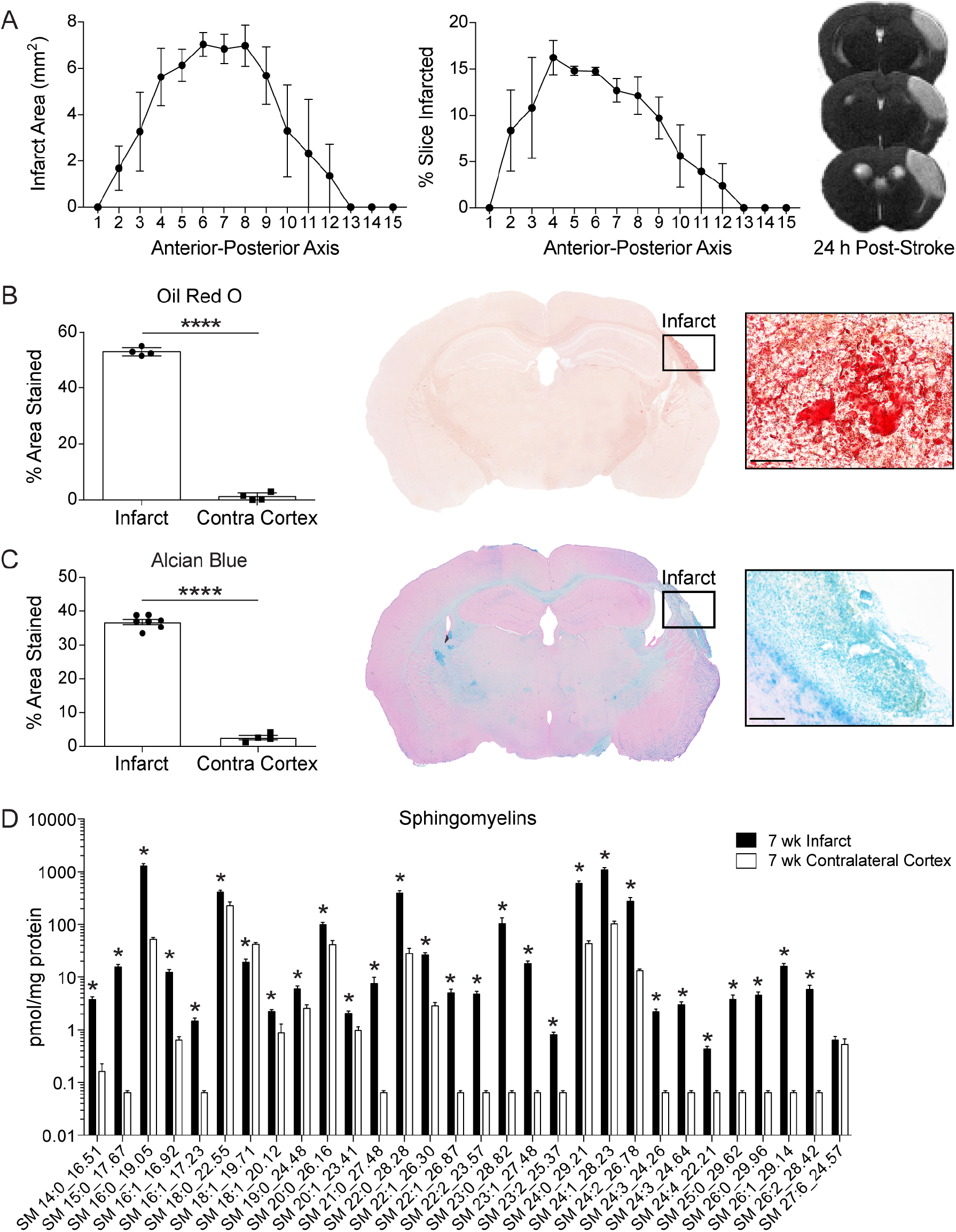
Lipid composition of chronic stroke infarcts in young adult mice. ***A*,** Infarct area per slice along the anterior-posterior axis determined by MRI at 24 h after DH stroke (*n* = 3; 0.8 mm slice thickness). *T_2_*-weighted images show infarcts (areas of hyperintense signal) centered on the somatosensory cortex, extending to the corpus callosum. ***B,*** Representative 2× and 60× images of Oil Red O staining show markedly more lipid droplets in infarcts than in contralateral cortices. Scale bar, 50 μm. (*n* = 4; paired *t* test; \*\*\*\**p* < 0.0001). ***C,*** Representative 2× and 20× images of Alcian blue staining show significantly more sulfatides in infarcts than in contralateral cortices. Scale bar, 125 μm. (*n* ≥ 4; unpaired *t* test; \*\*\*\**p* < 0.0001). ***D,*** Targeted lipidomics analysis revealed that infarcts had significantly higher levels of sphingomyelins than equivalent regions of the contralateral cortex at 7 weeks after stroke. (*n* = 8; multiple *t* tests, Holm– Sidak correction for multiple comparisons; \**p* < 0.05). Data are presented as mean ± SEM. SM, sphingomyelin.

We previously reported that, similar to atherogenesis, chronic stroke infarcts accumulate foamy macrophages, cholesterol crystals, and lipid droplets, which results in the upregulation of osteopontin, MMPs, and pro-inflammatory cytokines (Chung et al., 2018). To further evaluate the similarities between the pathophysiology of chronic stroke infarcts and atherosclerosis, we evaluated lipid droplet accumulation in chronic stroke infarcts. In the context of atherosclerosis, macrophages frequently become foamy in appearance due to the accumulation of cholesterol esters stored within lipid droplets. In addition to their role as storage containers within foamy macrophages, lipid droplets also contribute directly and indirectly to the pathology of progressing atherosclerotic plaques (Goldberg et al., 2018). To visualize lipid droplets in chronic stroke infarcts, we performed Oil Red O (ORO) staining on brain sections from young adult mice at 7 weeks after stroke. ORO staining revealed more intracellular lipid droplets in the infarcts compared to the contralateral cortices (**Fig. 1B**). These findings provide additional evidence that chronic stroke infarcts and atherosclerotic plaques share several similar pathophysiological features.

To assess whether lipid accumulation in chronic stroke infarcts is derived from myelin breakdown, we performed Alcian blue staining on brain sections from young adult mice at 7 weeks post-stroke. Alcian blue stains for sulfatide, a major constituent of CNS myelin. Importantly, sulfatide was recently identified as a myelin-associated inhibitor of neurite outgrowth (Winzeler et al., 2011). Chronic stroke infarcts contained an abundance of sulfatides at 7 weeks after stroke (**Fig. 1C**), indicating that phagocytic cells, including resident microglia and infiltrating macrophages, are strained in their capacity to process and eliminate myelin lipid debris for at least 7 weeks after stroke.

To further evaluate myelin-derived lipids in chronic stroke infarcts, we performed a targeted lipidomics analysis to assess sphingomyelin levels within infarcts compared to contralateral cortices of young adult mice at 7 weeks after stroke. Importantly, sphingomyelins play a central role in the structure of myelin, constituting 4% of myelin lipid content. The accumulation of sphingomyelin is a hallmark of Niemann–Pick disease and leads to changes in the plasma membrane that promote neurodegeneration (Mar et al., 2016). Sphingomyelins are also involved in signal transduction pathways and in the regulation of cholesterol and protein trafficking to myelin (Poitelon, Kopec, & Belin, 2020). In addition to their role in myelin architecture, sphingomyelins are involved in microglial activation and inflammation (Fitzner et al., 2020; Yang et al., 2020). Chronic stroke infarcts were enriched in sphingomyelins 7 weeks after stroke (**Fig. 1D**), indicating a pronounced dysregulation of myelin lipid homeostasis in chronic stroke infarcts of young adult mice.

### Lipid composition of chronic stroke infarcts in aged mice

Stroke remains the leading cause of long-term disability in people over the age of 65 (Virani et al., 2020). Therefore, we used aged (18-month-old) mice to further characterize the lipid composition of chronic stroke infarcts. First, to corroborate our observations in young adult mice, we used ORO to stain for lipid droplets in brain sections collected from aged mice at 7 weeks after stroke. We observed significantly more intracellular lipid droplets in infarcts compared to contralateral cortices (**Fig. 2A**). In addition, Alcian blue staining revealed that chronic stroke infarcts in aged mice are also characterized by an abundance of sulfatides when compared with contralateral cortices at 7 weeks after stroke (**Fig. 2B**). When comparing these results with those obtained in young adult mice, we see that aged mice have a comparable accumulation of lipid droplets and sulfatides at 7 weeks after stroke. Together, these results indicate that young adult and aged mice share similar impediments in the clearance of myelin lipid debris.

**Figure 2.**
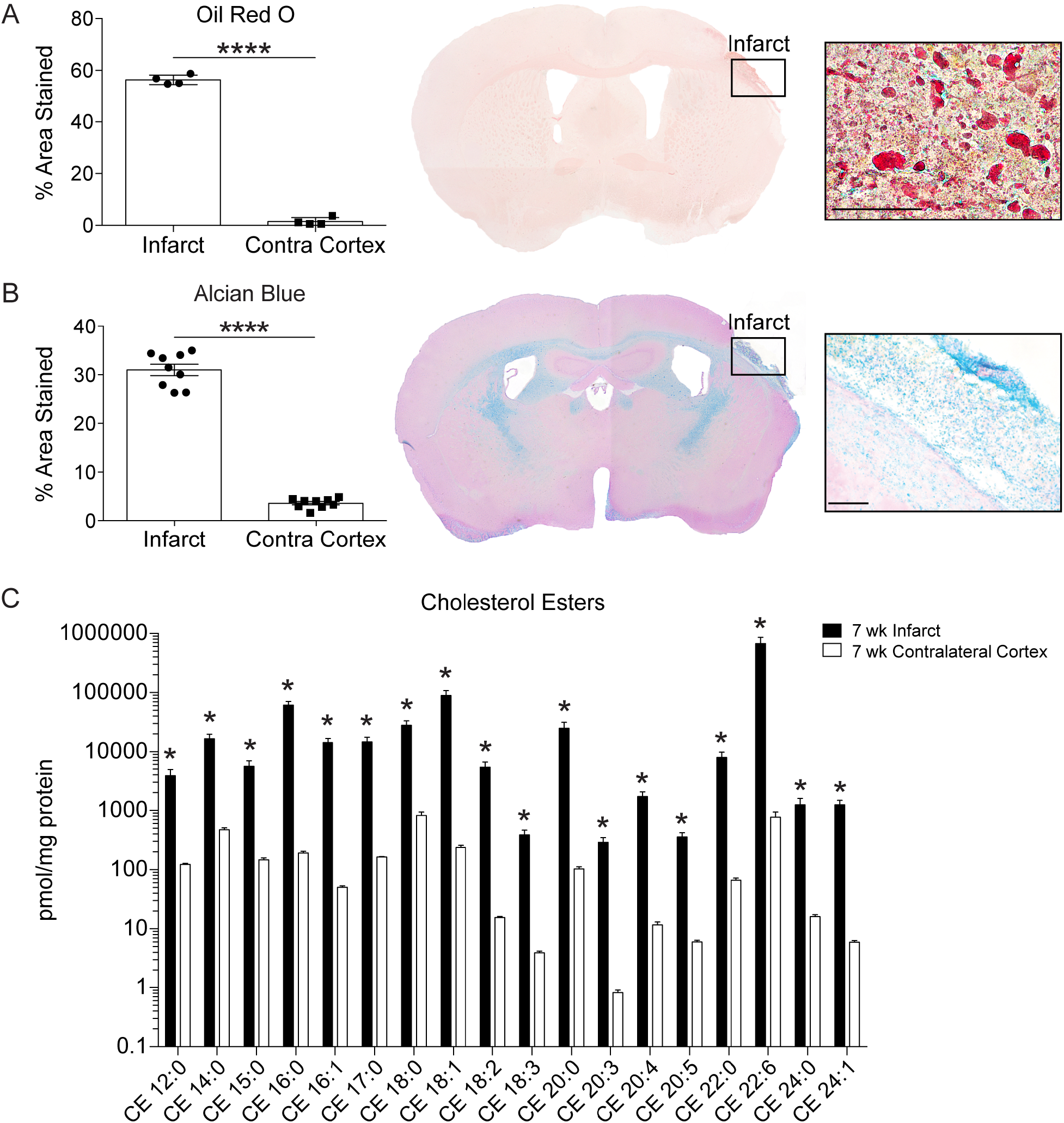
Lipid composition of chronic stroke infarcts in aged mice. ***A,*** Representative 2× and 60× images of Oil Red O staining show significantly more lipid droplets in infarcts compared to contralateral cortices. Scale bar, 50 μm. (*n* = 4; paired *t* tests; \*\*\*\**p* < 0.0001). ***B,*** Representative 2× and 20× images of Alcian blue staining show significantly more sulfatides in infarcts compared to contralateral cortices. Scale bar, 125 μm. (*n* ≥ 4; paired *t* tests; \*\*\*\**p* < 0.0001). ***C,*** Targeted lipidomics analysis revealed significantly higher levels of cholesterol esters in infarcts than in equivalent regions of the contralateral cortex at 7 weeks after stroke. (*n* = 5; multiple *t* tests, Holm– Sidak correction for multiple comparisons; \**p* < 0.05). Data are presented as mean ± SEM. CE, cholesterol ester.

Cholesterol is esterified prior to storage in cytoplasmic lipid droplets to prevent free cholesterol-associated cell toxicity (Ghosh et al., 2010). Therefore, we evaluated the composition of cholesterol esters stored in lipid droplets via a second targeted lipidomics analysis. For this analysis, we again compared infarcts at 7 weeks after stroke to equivalent regions of the contralateral cortex. All quantified cholesterol ester species were significantly elevated in infarcts of aged mice at 7 weeks after stroke compared to contralateral cortices (**Fig. 2C**).

### Transcriptome of the chronic stroke infarct in young adult mice

To systematically characterize the chronic stroke infarct transcriptome, we conducted bulk RNA sequencing (RNA-Seq) on infarcts collected at 7 weeks after stroke and compared them to equivalent regions of the contralateral cortex (**Fig. 3A**). As expected, differential expression (DE) analysis revealed marked differences between infarcts and contralateral cortices. The resultant volcano plot illustrates the distribution of 3,930 upregulated and 2,869 downregulated genes (**Fig. 3B**). The most significantly upregulated genes, including *Lpl*, *Spp1*, *Cd36*, *Mmp2*, and *Mmp19*, and the most highly enriched genes, including *Cd5l*, *Mmp3*, *Mmp12*, and *Mmp13*, indicate a pronounced disturbance in lipid homeostasis in infarcts at 7 weeks after stroke. Importantly, Zhu et al. (2017) similarly identified *Cd5l*, *Lpl*, and *Cd36* as highly expressed genes in foamy macrophages after spinal cord injury (SCI) (Zhu et al., 2017).

**Figure 3.**
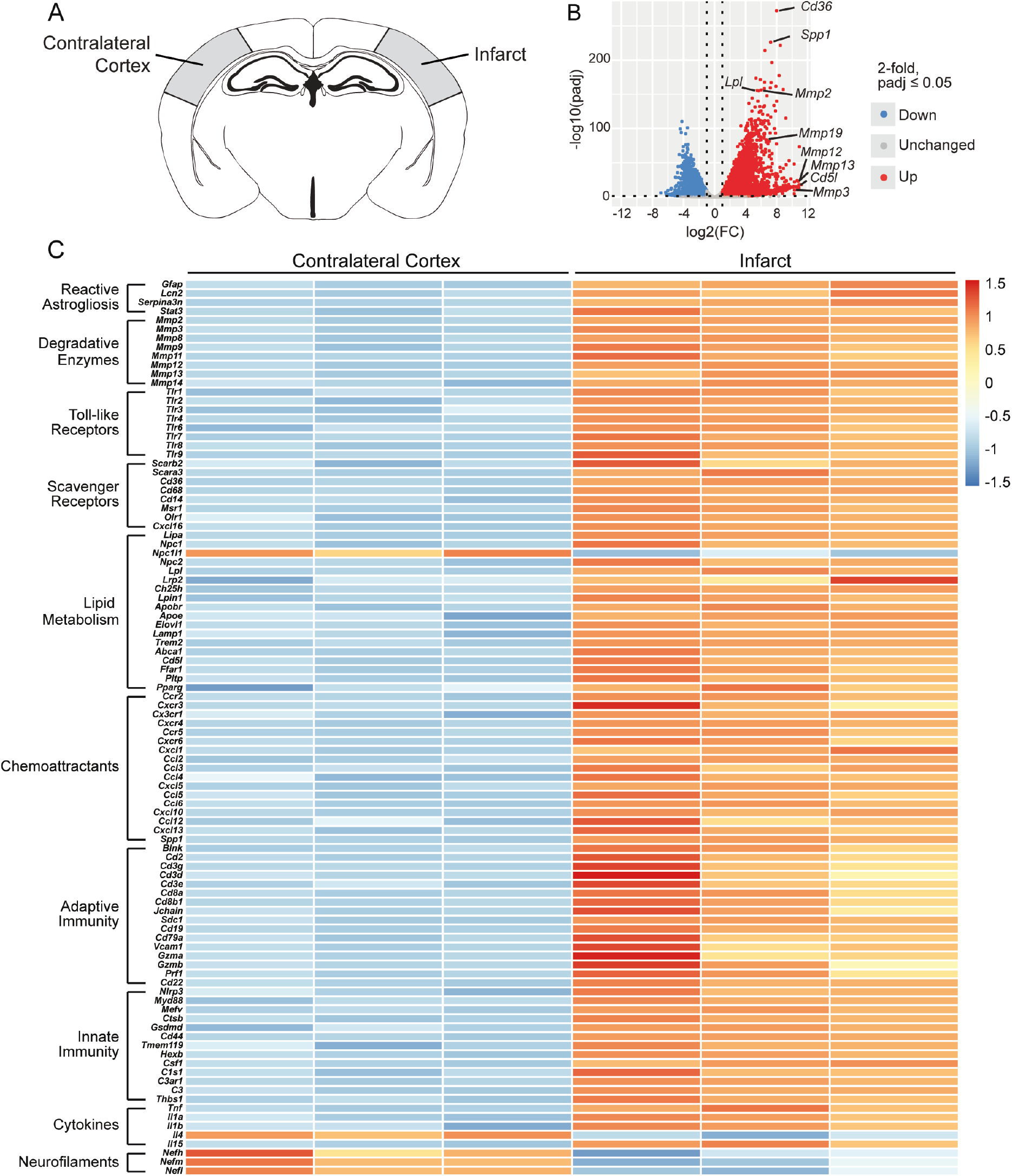
Transcriptome of the chronic stroke infarct in young adult mice. ***A,*** Schematic of a mouse coronal brain section following stroke induced by distal middle cerebral artery occlusion + hypoxia, with the analyzed region of each hemisphere shaded in gray. ***B,*** Volcano plot showing differences in gene expression between the infarct and contralateral cortex at 7 weeks following stroke (false discovery rate- adjusted *p* < 0.05; FC > |2|). ***C,*** Row-scaled heatmap displaying differentially expressed genes associated with dysregulated lipid metabolism and pronounced inflammation (false discovery rate-adjusted *p* < 0.05; FC > |2|). FC, fold change.

Correspondingly, chronic stroke infarcts were also characterized by an upregulation of TLRs, including *Tlr1*, *Tlr2*, and *Tlr4*, and scavenger receptors, including *Scarb2*, *Cd68*, *Msr1*, and *Cxcl16*. These genes are implicated in the induction and progression of atherosclerosis; importantly, these scavenger receptors mediate the uptake of oxLDL into macrophages and promote their differentiation into foam cells (Aslanian & Charo, 2006; Febbraio, Hajjar, & Silverstein, 2001; Park, 2014; Rahaman et al., 2006; Zeibig et al., 2019). Dysregulated lipid homeostasis within infarcts is further evidenced by the upregulation of genes involved in lipid metabolism: *Npc1*, *Npc2*, *Lamp1*, *Ffar1*, *Pparg*, *Apoe*, *Trem2*, and *Abca1* (**Fig. 3C**).

In addition to alterations in lipid metabolism, the transcriptome also revealed a signature of chronic inflammation at 7 weeks after stroke. For instance, activation of the innate immune response is indicated by the upregulation of genes involved in complement cascade activation, including *C1s1*, *C3ar1*, and *C3*, and in phagocytosis, including *Tmem119*, *Hexb*, *Nlrp3*, *Ctsb*, and *Myd88*. Further, upregulated genes included *Gfap*, *Lcn2*, and *Serpina3n*, which have been validated as specific markers of reactive astrogliosis in ischemic stroke and LPS-induced neuroinflammation (Zamanian et al., 2012) (**Fig. 3C**). Together, these genes signify the activation of innate immune cells, including infiltrating macrophages, resident microglia, and astrocytes.

Additionally, activation of the adaptive immune system is indicated by the upregulation of genes associated with T lymphocytes, including *Cd2*, *Cd3e*, *Gzma*, *Gzmb*, and *Prf1*; B lymphocytes, including *CD19*, *Cd79a*, and *Blnk*; and antibody-producing plasma cells, including *Sdc1* and *Jchain* (**Fig. 3C**). These perturbations in innate and adaptive immunity demonstrate that chronic inflammation persists in infarcts for at least 7 weeks after stroke, as we have shown previously (Chung et al., 2018; Doyle et al., 2015; Zbesko et al., 2021).

In conjunction with the upregulation of genes involved in chronic inflammation and lipid metabolism, the DE analysis revealed a downregulation of genes involved in neuronal structure and function. Specifically, the neurofilament subunits *Nefh*, *Nefm*, and *Nefl* were downregulated, which indicates persistent disruption or absence of neuronal cytoskeletons in infarcts at 7 weeks after stroke (**Fig. 3C**).

We then used IPA software to define upstream regulators and identify altered biological processes based on DE analysis. Differentially expressed genes were associated with the following upregulated biological processes: (i) atherosclerosis signaling, (ii) IL-1 signaling, (iii) inflammasome pathway, (iv) eicosanoid signaling, (v) B cell receptor (BCR) signaling, and (vi) phospholipases, along with others. In contrast, differentially expressed genes were associated with the following downregulated biological processes: (i) calcium signaling, (ii) glutamate receptor signaling, and (iii) synaptic long-term potentiation (**Fig. 4A, B**). These altered biological processes indicate that the stroke infarct transcriptome is characterized by chronic inflammation, dysregulated lipid metabolism, and impaired or absent neuronal function at 7 weeks after stroke. In addition, IPA identified lipid metabolic upstream regulators such as cholesterol, phospholipids, and LDL, as well as immunological upstream regulators such as MYD88, IL18, CD3, IL1B, TLR4, and TNF (**Fig. 4C**).

**Figure 4.**
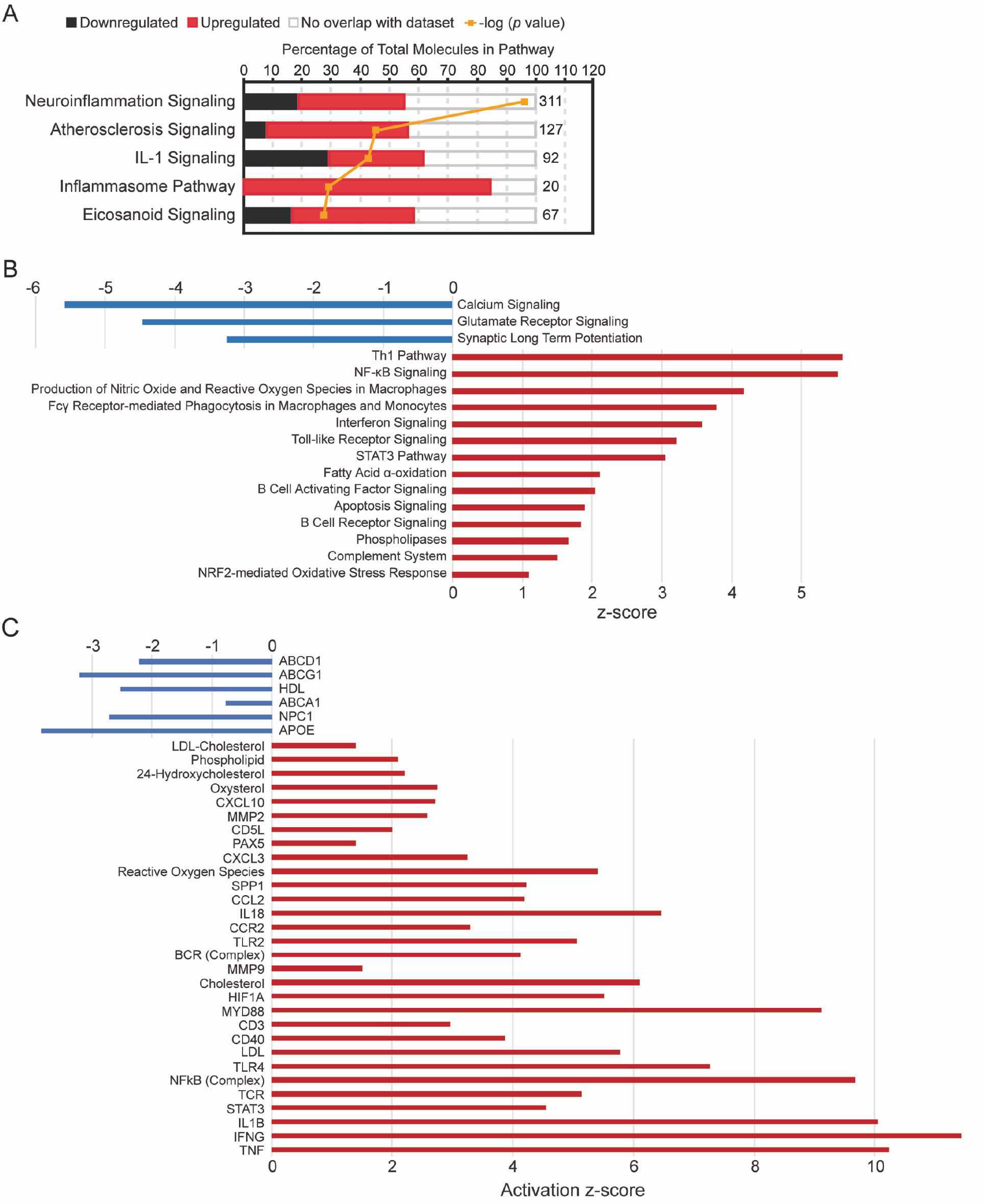
Canonical pathways and upstream regulators based on differentially expressed genes in chronic stroke infarcts of young adult mice. ***A,*** Ingenuity Pathway Analysis (IPA) performed on differentially expressed genes revealed a significant upregulation of inflammatory pathways in infarcts of young adult mice at 7 weeks after stroke compared to contralateral cortices. ***B,*** Additional canonical pathway analysis confirmed activation of inflammatory pathways in infarcts of young adult mice at 7 weeks after stroke. ***C,*** IPA identified putative upstream regulators associated with inflammation and lipid metabolism, including cholesterol, LDL, MYD88, CD3, and IL1B. All canonical pathways and upstream regulators shown are significantly up- or downregulated (*p* < 0.05).

### HPβCD attenuates the chronic inflammatory response to stroke in young adult mice

To investigate the role of lipid metabolism in the chronic inflammatory response to stroke, we administered subcutaneous injections of 4 g/kg HPβCD or vehicle control to young adult mice triweekly (Monday, Wednesday, and Friday) for 6 weeks, beginning 1 week after stroke (**Fig. 5A**). We performed immunostaining on vehicle- and HPβCD- treated brain sections at 7 weeks post-stroke. These analyses revealed substantially fewer B220+ B lymphocytes, CD3ε+ T lymphocytes, and CD138+ and IgA+ antibody-producing plasma cells in infarcts of HPβCD-treated mice than in infarcts of vehicle-injected mice. In addition, ORO staining revealed that infarcts of HPβCD-treated mice had fewer lipid droplets than those of vehicle-injected mice (**Fig. 5B**). These differences indicate that repeated administration of HPβCD following stroke attenuates chronic inflammation and lipid droplet formation in young adult mice. Conversely, the accumulation of sulfatides in infarcts of vehicle- and HPβCD-treated mice was comparable (**Fig. 5B**), suggesting that HPβCD does not aid in the clearance of sulfatides.

**Figure 5.**
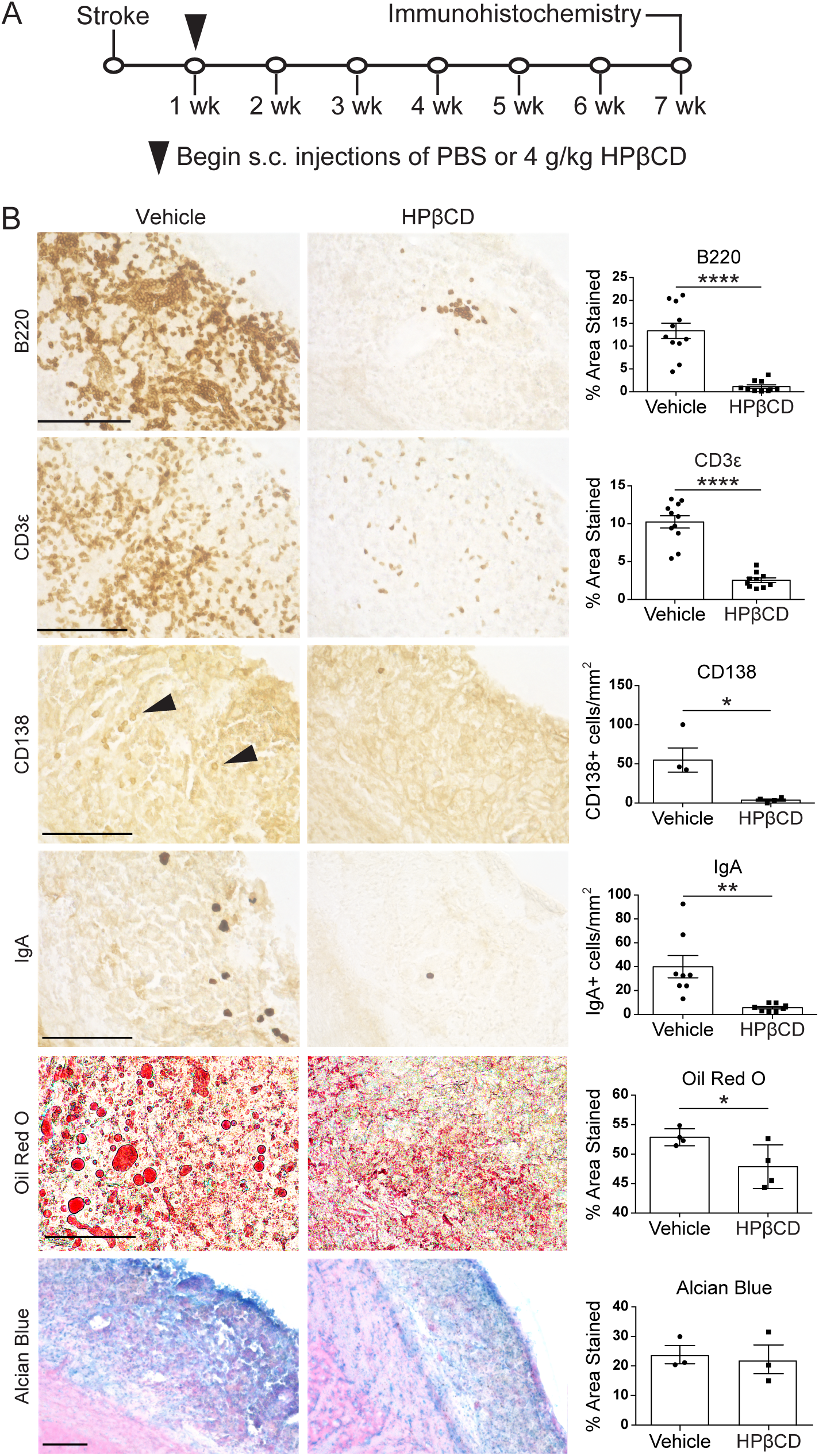
HPβCD attenuates the chronic inflammatory response to stroke in young adult mice. ***A,*** Experimental design, *n* = 31–32 per group. Three-month-old mice received subcutaneous (s.c.) injections of 4 g/kg HPβCD or vehicle three times a week, beginning 1 week after stroke induced by distal middle cerebral artery occlusion + hypoxia. Brains were extracted at 7 weeks post-stroke and processed for immunohistochemistry. ***B,*** Representative 40× images of infarcts in vehicle- or HPβCD- treated young adult mice stained for B lymphocytes (B220), T lymphocytes (CD3ε), and antibody-producing plasma cells (CD138, IgA), followed by representative 60× images of infarcts stained for lipid droplets (Oil Red O) and 20× images of infarcts stained for sulfatides (Alcian blue). Quantification of images is shown to the right of each photomicrograph. (*n* = 3–11; unpaired *t* tests; \**p* < 0.05, \*\**p* < 0.01, \*\*\*\**p* < 0.0001). Scale bars, 50 μm (Oil Red O), 125 μm (B220, CD3ε, CD138, IgA, Alcian blue). Data are presented as mean ± SEM.

### HPβCD attenuates the chronic inflammatory response to stroke in aged mice

Stroke prevalence and mortality rates increase with advancing age in both males and females (Virani et al., 2020). Therefore, to address age as a biological variable, we performed immunohistochemistry and bulk RNA-Seq analyses on aged (18-month-old) mice treated with HPβCD using the same treatment regimen as in young adult mice (HPβCD or vehicle control three times per week for 6 weeks) (**Fig. 6A**). We discovered that, consistent with young adult mice, aged mice had substantially fewer B220+ B lymphocytes, CD3ε+ T lymphocytes, and IgA+ antibody-producing plasma cells in their infarcts following repeated administration of HPβCD. In addition, ORO staining revealed that infarcts of HPβCD-treated mice had significantly less lipid droplet accumulation than vehicle-injected mice at 7 weeks after stroke (**Fig. 6B**). These results demonstrate that young adult and aged mice similarly exhibit attenuated lipid droplet and immune cell accumulation in chronic stroke infarcts following repeated administration of HPβCD. However, the accumulation of sulfatides in vehicle- and HPβCD-treated aged mice was comparable (**Fig. 6B**), reinforcing that HPβCD does not aid in the clearance of sulfatides.

**Figure 6.**
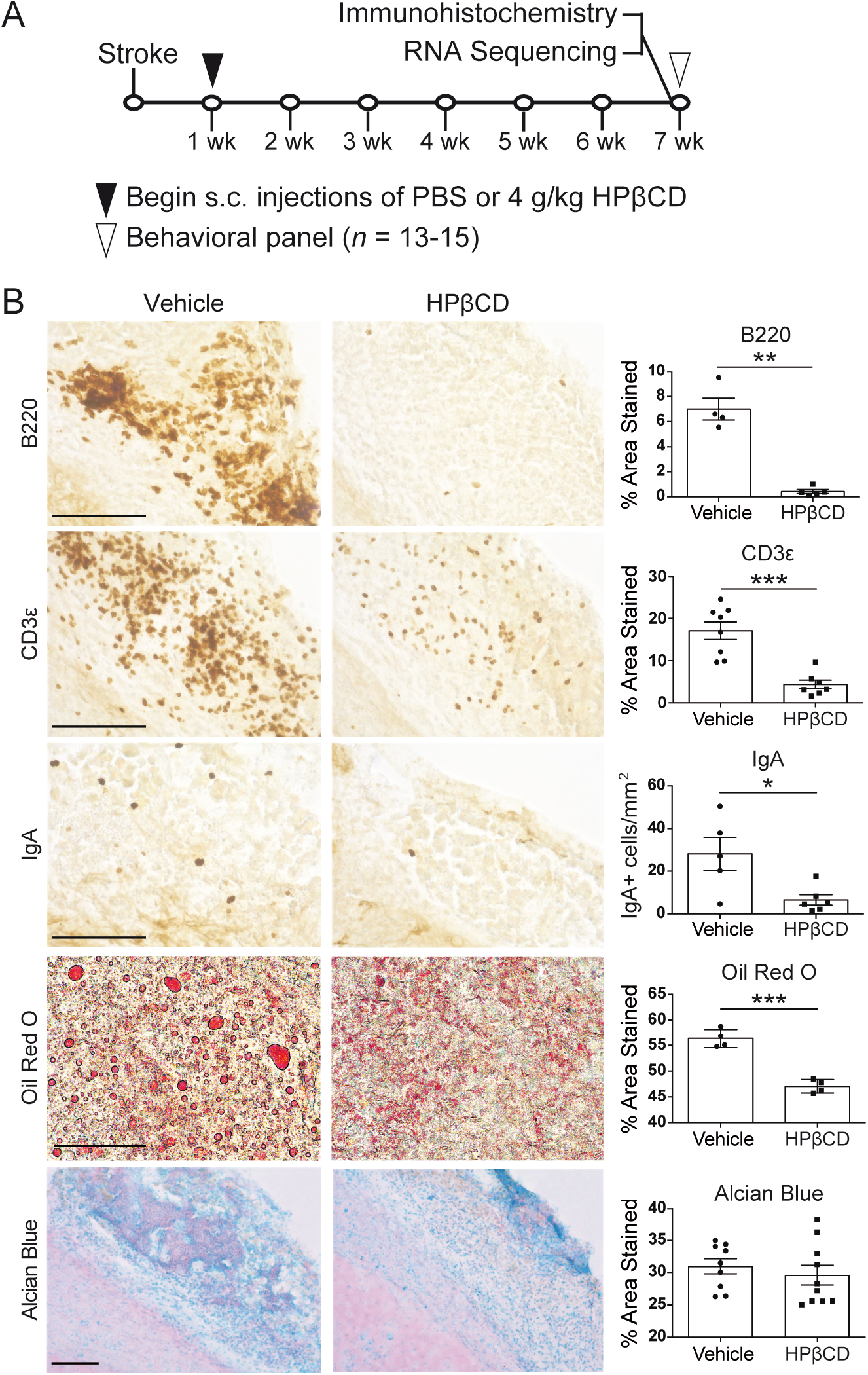
HPβCD attenuates the chronic inflammatory response to stroke in aged mice. ***A,*** Experimental design, *n* = 39–40 per group. Eighteen-month-old mice received subcutaneous (s.c.) injections of 4 g/kg HPβCD or vehicle three times a week, beginning 1 week after stroke induced by distal middle cerebral artery occlusion + hypoxia. Brains were dissected at 7 weeks post-stroke and processed for immunohistochemistry or RNA-Seq. ***B,*** Representative 40× images of infarcts in vehicle- or HPβCD-treated young adult mice stained for B lymphocytes (B220), T lymphocytes (CD3ε), and antibody-producing plasma cells (IgA), followed by representative 60× images of infarcts stained for lipid droplets (Oil Red O) and 20× images of infarcts stained for sulfatides (Alcian blue).Quantification of images is shown to the right of each photomicrograph. (*n* = 4–10; unpaired *t* tests; \**p* < 0.05, \*\**p* < 0.01, \*\*\**p* < 0.001). Scale bars, 50 μm (Oil Red O), 125 μm (B220, CD3ε, IgA, Alcian blue). Data are presented as mean ± SEM.

### HPβCD does not alter peripheral immune cell populations in the blood or spleens of aged mice after stroke

The spleen, a secondary lymphoid organ, is a major reservoir of immune cells and a focal point for the immune response to tissue injury. In response to ischemic stroke, splenocytes enter into systemic circulation and migrate to the brain, exacerbating neurodegeneration (Seifert et al., 2012). To determine the effect of HPβCD on peripheral immune cell populations, we used flow cytometry to quantify splenic cell populations. There were no significant differences in CD19+ B lymphocytes or GL-7+ germinal center B lymphocytes (**Fig. 7A, B**), or in IgG1 and IgA isotypes (**Fig. 7C**), between the spleens of vehicle- and HPβCD-treated mice at 7 weeks after stroke. The CD4+ T lymphocyte, CD8+ T lymphocyte, and Treg populations were also not significantly altered by repeated HPβCD administration (**Fig. 7D, E**). These results demonstrate that splenic cell populations in HPβCD-treated aged mice are not significantly altered in comparison to those of vehicle-injected mice.

**Figure 7.**
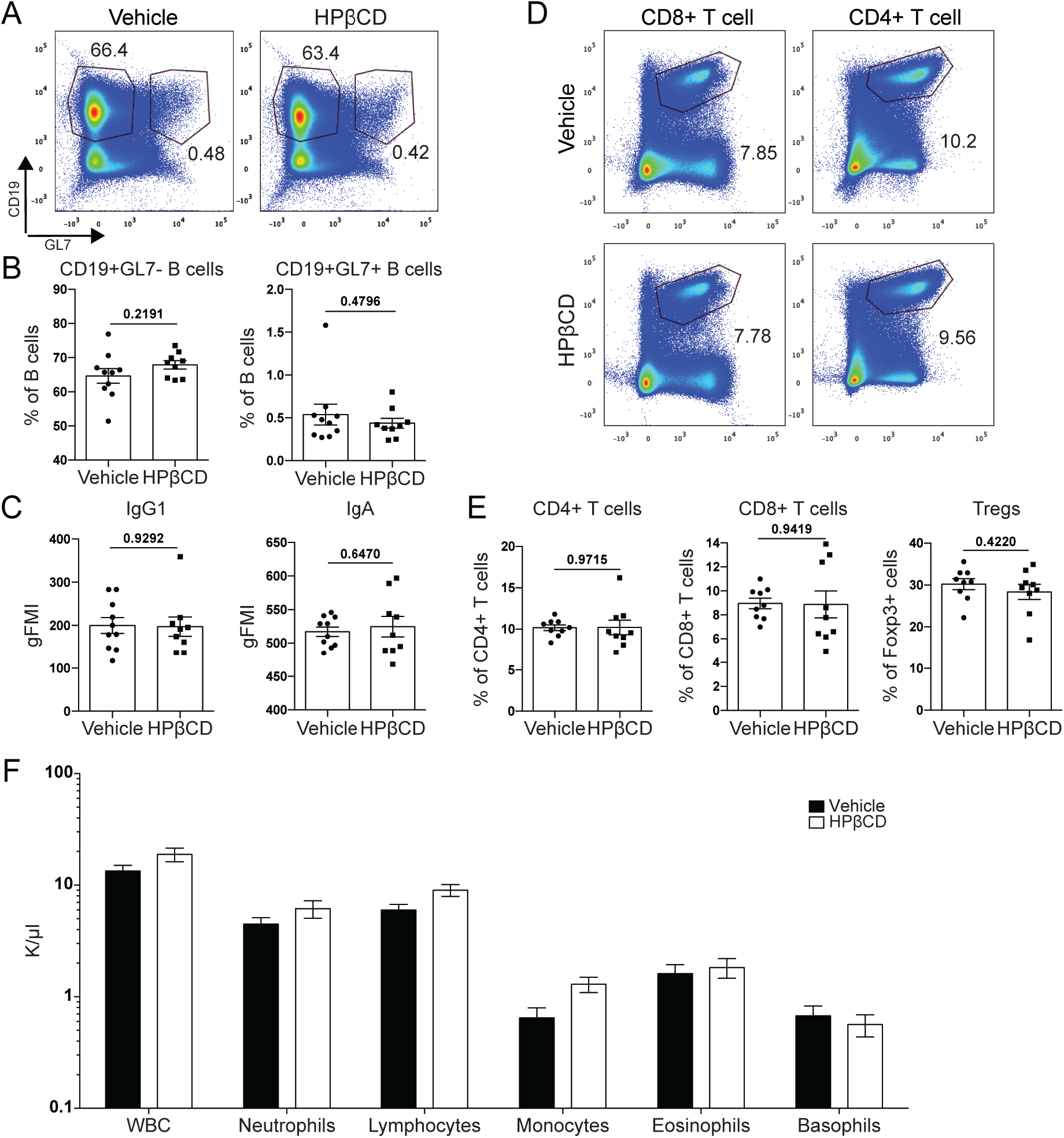
HPβCD does not alter peripheral immune cell populations in the blood or spleens of aged mice after stroke. ***A, B,*** Splenocytes from HPβCD- and vehicle-injected aged mice were stained with antibodies against CD19 and GL7. Representative flow cytometry plots (***A***) and quantification of CD19+GL7- B cells and CD19+GL7+ germinal center B cells as a percentage of total lymphocytes (***B***) showed no significant differences in splenic B cell populations between vehicle- and HPβCD-treated aged mice at 7 weeks after stroke. ***C,*** Quantification of antibody isotypes (IgG1 and IgA) in the spleen revealed no significant differences between vehicle- and HPβCD-treated aged mice at 7 weeks after stroke. ***D, E,*** Splenocytes from HPβCD- and vehicle-injected aged mice were stained with antibodies against TCRβ, CD4, CD8, and Foxp3. Representative flow cytometry plots of CD4+ and CD8+ T cells (***D***) and quantification of CD4+ T cells, CD8+ T cells, and Tregs as a percentage of total lymphocytes (for CD4+ and CD8+ gates) or TCRβ+CD4+ T cells (for Treg gate) (***E***) revealed no significant differences in splenic T cell populations between vehicle- and HPβCD-treated aged mice at 7 weeks after stroke. (*n* = 9**–**10; unpaired *t* tests; \**p* < 0.05). ***F,*** Complete blood count analysis in vehicle- and HPβCD-treated aged mice at 7 weeks after stroke revealed no significant differences in circulating immune cell populations (*n* = 6; multiple *t* tests, Holm–Sidak correction for multiple comparisons; \**p* < 0.05). Data are presented as mean ± SEM. GC, germinal center.

A complete blood count analysis on blood collected from vehicle- and HPβCD-treated aged mice at 7 weeks after stroke also revealed no significant differences in the number of circulating white blood cells, neutrophils, lymphocytes, monocytes, eosinophils, or basophils (**Fig. 7F**). Together, the flow cytometry and complete blood count analyses indicate that, while there were substantially fewer immune cells in infarcts of HPβCD- treated aged mice than in those of vehicle-injected mice at 7 weeks after stroke (**Fig. 6B**), repeated systemic HPβCD administration does not significantly alter circulating or splenic immune cell populations in aged mice.

### HPβCD promotes lipid metabolism and attenuates inflammation in infarcts of aged mice after stroke

We systematically characterized the transcriptomes of stroke infarcts in vehicle- and HPβCD-treated aged mice by performing bulk RNA-Seq on infarcted brain tissue collected 7 weeks after stroke (**Fig. 8A**). As shown in the volcano and MA plots, the DE analysis revealed 991 upregulated genes and 1,876 downregulated genes (**Fig. 8B, C**). HPβCD treatment upregulated intracellular cholesterol transporters, lipoproteins, and lipoprotein receptors, such as *Npc1*, *Npc2*, *Apoe*, *Trem2*, *Abca1*, *Vldlr*, and *Hdlbp*, and downregulated *Ffar1*, *Lrpb1*, and *Dgkb* (**Fig. 8D, E**). These alterations in lipid metabolism suggest that HPβCD aids in the restoration of lipid homeostasis by 7 weeks after stroke. In addition, the DE analysis revealed a downregulation of genes involved in chemotaxis, including *Cx3cr1*, *Cxcl5*, and *Ccr6*, and innate and adaptive immune responses, including *Mmd2*, *Mzb1*, *Btla*, *Jchain*, and *Igkv5-48*. Correspondingly, there was an upregulation of anti-inflammatory genes, such as *Tgfb1*, *Tgfb2*, *P2ry2*, and *Il18bp* (**Fig. 8D**). These results indicate that HPβCD attenuates the chronic inflammatory response to stroke in aged mice.

**Figure 8.**
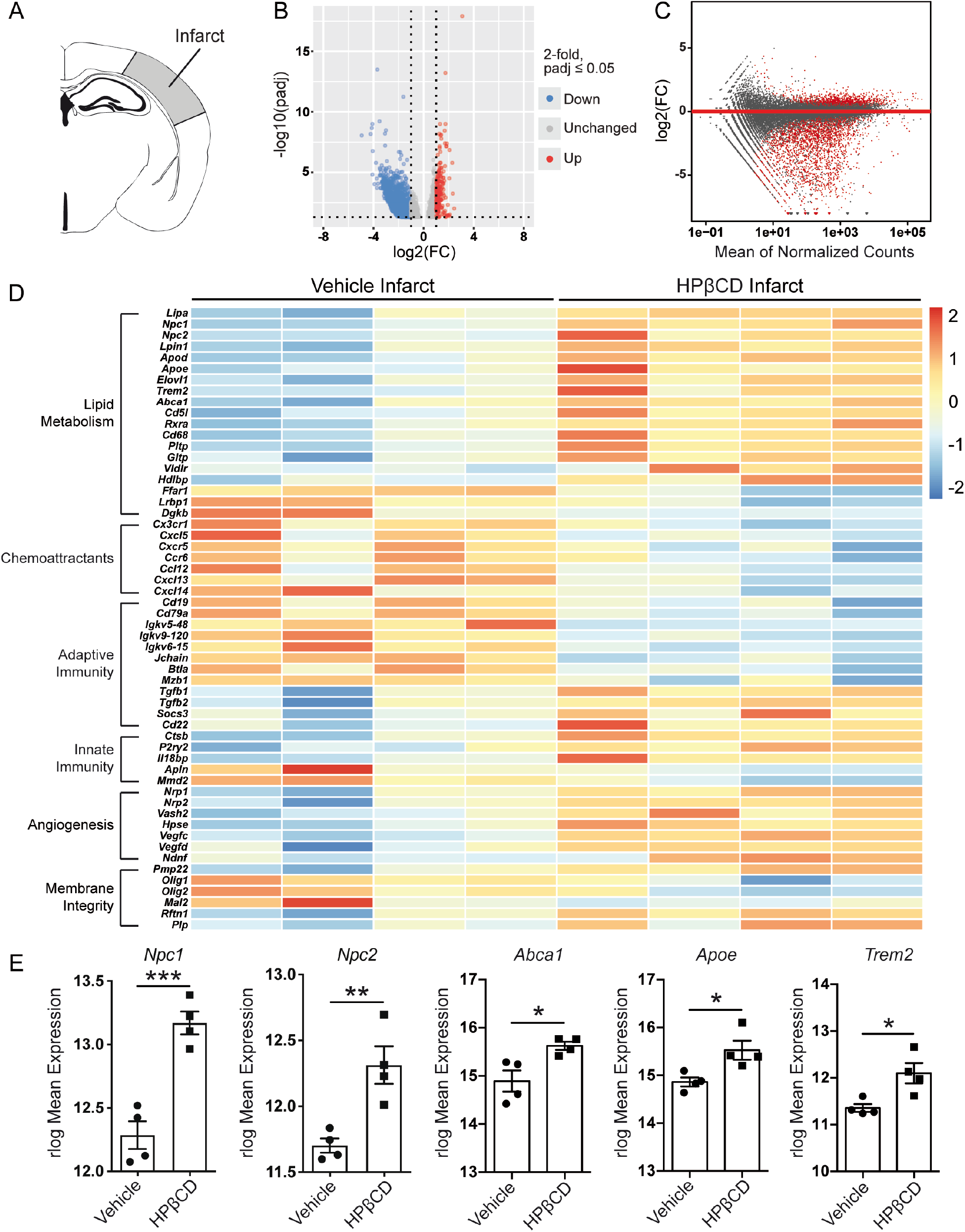
HPβCD promotes lipid metabolism and attenuates inflammation in infarcts of aged mice after stroke. ***A,*** Schematic of a mouse coronal brain section following stroke induced by distal middle cerebral artery occlusion + hypoxia, with the analyzed region shaded in gray. ***B, C,*** Volcano (***B***) and MA (***C***) plots constructed from count data show differences in gene expression between infarcts from vehicle- and HPβCD-treated mice at 7 weeks after stroke (false discovery rate-adjusted *p* < 0.05; FC > |2|). ***D,*** Row-scaled heatmap displaying differentially expressed genes associated with lipid metabolism, inflammation, and angiogenesis (false discovery rate-adjusted *p* < 0.05; FC > |2|). ***E,*** Graphs representing rlog-normalized expression of selected genes related to lipid metabolism. (*n* = 4; unpaired *t* tests; \**p* < 0.05, \*\**p* < 0.01, \*\*\**p* < 0.001). FC, fold change.

We also identified alterations in genes involved in the composition of membranes, including *Olig1*, *Pmp22*, *Mal2*, *Rftn1*, and *Plp*. These alterations suggest that HPβCD modifies lipid rafts and cell membranes by 7 weeks after stroke. Genes involved in angiogenesis were also upregulated, including *Nrp1*, *Vash2*, *Hpse*, *Vegfc*, *Vegfd*, and *Ndnf* (**Fig. 8D**). In addition, enrichment maps constructed from GO terms revealed an upregulation of lipid metabolism, angiogenesis, and collagen synthesis pathways and a downregulation of immune response pathways (**Fig. 9A**). These results indicate that HPβCD treatment promotes lipid metabolism and angiogenesis and dampens chronic inflammation in infarcts of aged mice at 7 weeks after stroke.

**Figure 9.**
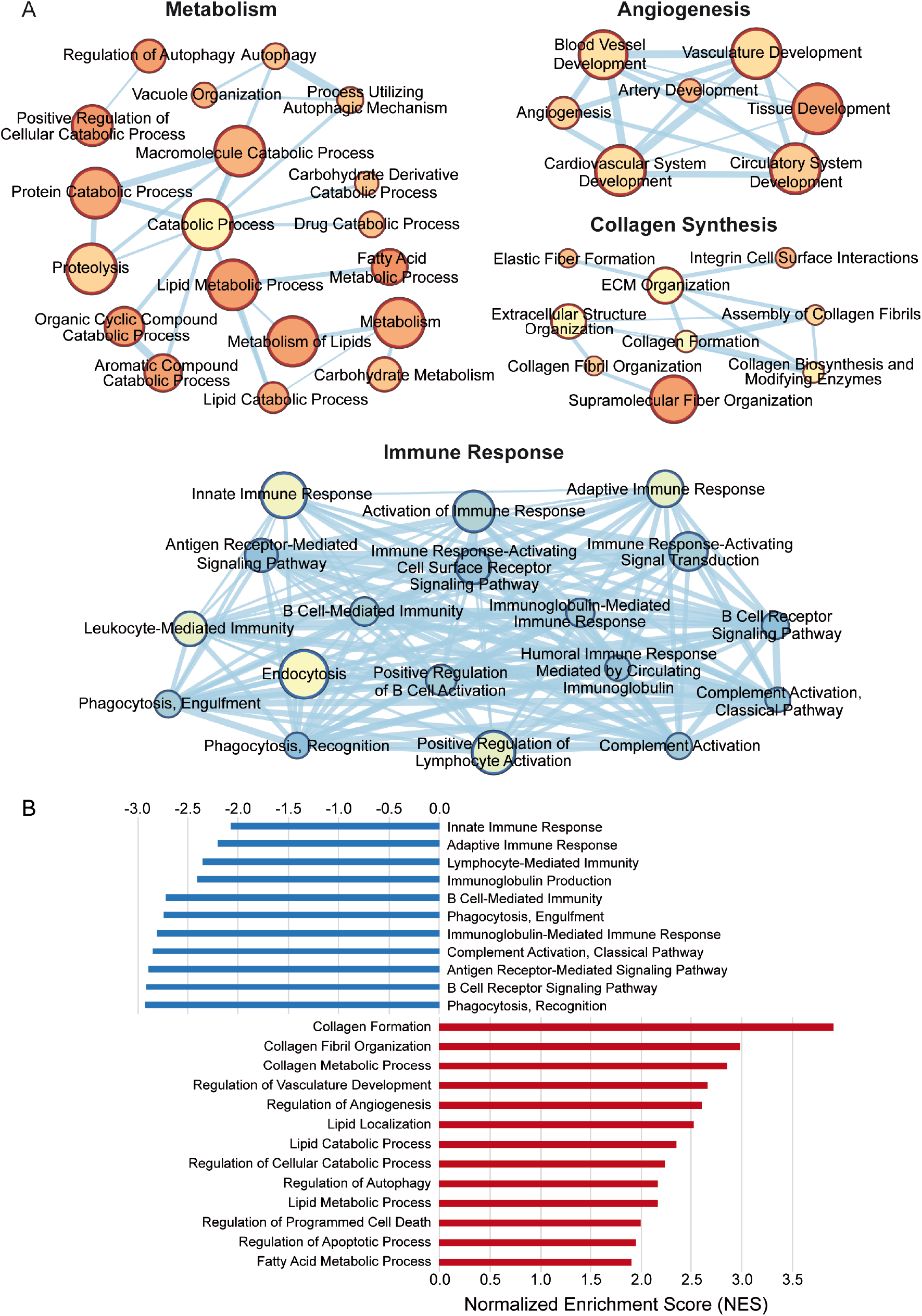
Biological processes enriched in infarcts of HPβCD-treated aged mice at 7 weeks after stroke. ***A,*** Enrichment maps constructed from Gene Ontology terms revealed a significant upregulation of metabolism, angiogenesis, and collagen synthesis pathways and a significant downregulation of immune response pathways. Pathways are shown as circles (nodes) that are connected with lines (edges) if the pathways share genes. Node colors are based on the enrichment score, and edge sizes are based on the number of genes shared by the connected pathways. ***B,*** Gene set enrichment analysis (GSEA) revealed significant enrichment of various biological processes in infarcts of HPβCD-treated aged mice, depicted in red. Conversely, GSEA identified biological processes more significantly enriched in infarcts of vehicle-injected mice compared to HPβCD-treated mice, depicted in blue.

Next, we performed genome-wide expression analysis of the RNA-Seq data using GSEA, which revealed significant enrichment of various biological processes in infarcts of HPβCD-treated aged mice, including (i) collagen formation, (ii) regulation of angiogenesis, (iii) lipid catabolic process, (iv) regulation of autophagy, and (v) regulation of apoptotic process. Conversely, GSEA identified biological processes more enriched in infarcts of vehicle-injected mice compared to HPβCD-treated mice, including (i) innate immune response, (ii) adaptive immune response, (iii) immunoglobulin production, (iv) phagocytosis, engulfment, and (v) complement activation, classical pathway (**Fig. 9B**). These results further indicate that infarcts of vehicle-injected mice are defined by increased innate and adaptive immune responses, whereas those of HPβCD-treated mice are defined by increased angiogenic and lipid metabolic processes.

### Transcriptome of the peri-infarct region in aged mice at 7 weeks after stroke

Prior to assessing the impact of repeated HPβCD administration on the peri-infarct region of the brain, we first compared the transcriptome of the peri-infarct region to the equivalent region of the contralateral hemisphere in aged mice at 7 weeks after stroke (**Fig. 10A**). The peri-infarct region was characterized by 3,910 differentially expressed genes (**Fig. 10B, C**). Of these, 2,123 were upregulated, many of which were also upregulated in infarcts at 7 weeks after stroke (**Fig. 3C**). For instance, the infarct and peri-infarct regions were both defined by the upregulation of degradative enzymes, including *Mmp2*, *Mmp3*, and *Mmp8*; TLRs, including *Tlr1*, *Tlr2*, and *Tlr4*; and scavenger receptors, including *Cd36*, *Cd68*, and *Cxcl16*. Additionally, peri-infarct regions were characterized by specific markers of reactive astrogliosis, including *Gfap*, *Lcn2*, and *Serpina3n* (**Fig. 10D**).

**Figure 10.**
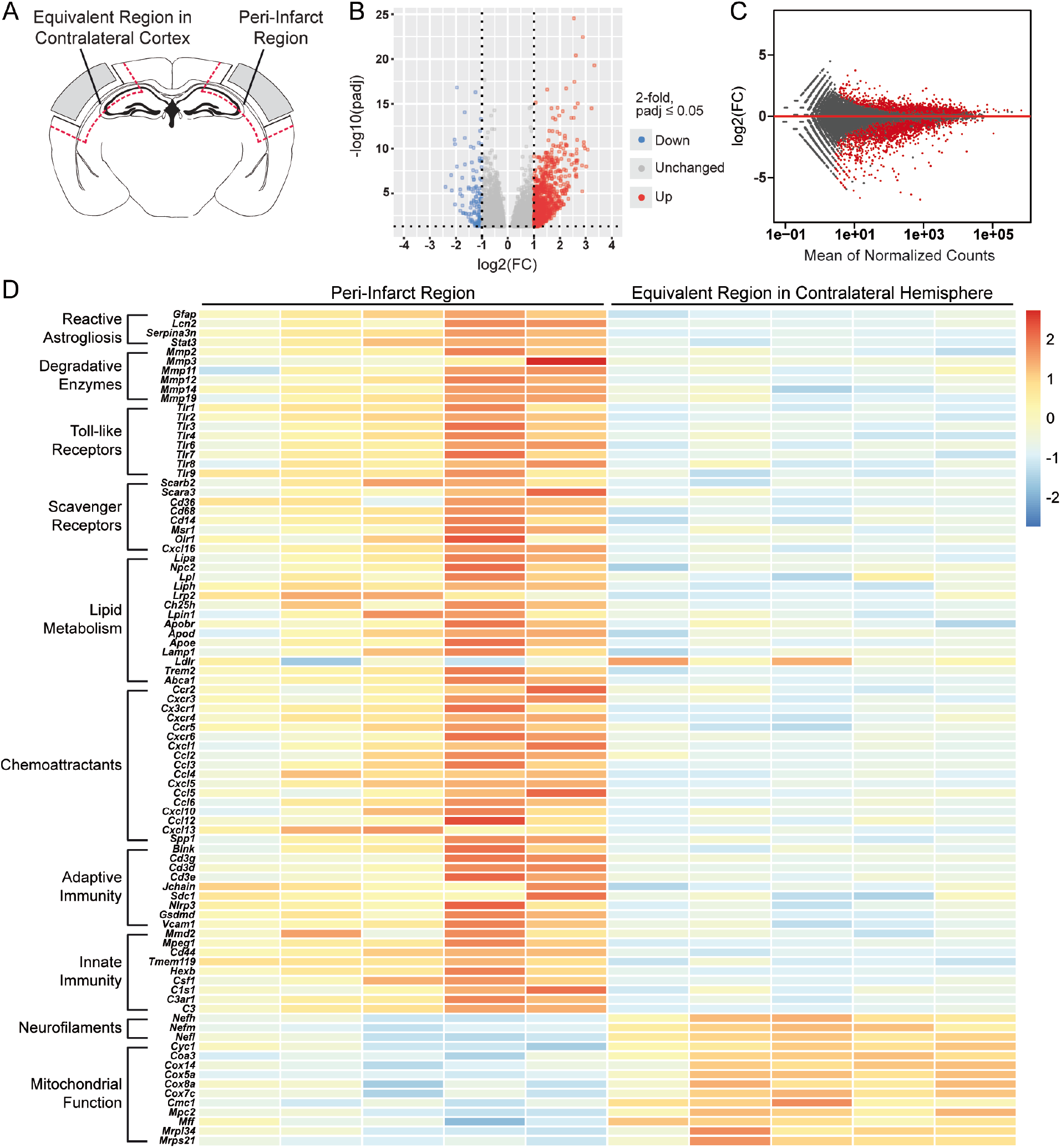
Transcriptome of the peri-infarct region in aged mice at 7 weeks after stroke. ***A,*** Schematic of a mouse coronal brain section following stroke induced by distal middle cerebral artery occlusion + hypoxia, with analyzed regions indicated. ***B, C,*** Volcano (***B***) and MA (***C***) plots constructed from count data show differences in gene expression between peri-infarct regions and equivalent regions in the contralateral cortex at 7 weeks after stroke (false discovery rate-adjusted *p* < 0.05; FC > |2|). ***D,*** Row- scaled heatmap displaying differentially expressed genes associated with lipid metabolism, inflammation, and mitochondrial function (false discovery rate-adjusted *p* < 0.05; FC > |2|). FC, fold change.

Similar to the transcriptome of the infarct at 7 weeks after stroke, the transcriptome of the peri-infarct region was primarily defined by genes involved in chronic inflammation and dysregulated lipid metabolism. The upregulation of *Lpl*, *Lrp2*, *Ch25h*, *Apoe*, *Trem2*, and *Abca1* signifies a divergence from lipid homeostasis. Moreover, *Ldlr* was downregulated, which implies impaired lipoprotein and lipid metabolism (Go & Mani, 2012). In addition, genes involved in complement cascade activation (including *C1s1*, *C3ar1*, and *C3*) and phagocytosis (including *Tmem119*, *Hexb*, *Nlrp3*, and *Mmd2*) were upregulated in peri-infarct regions. These alterations in innate immunity coincided with corresponding perturbations in adaptive immunity. The upregulation of *Cd3d*, *Cd3e*, *Cd3g*, *Blnk*, *Sdc1* and *Jchain* suggests that chronic inflammation persists in peri-infarct regions for at least 7 weeks after stroke (**Fig. 10D**).

Conversely, the DE analysis revealed that 1,787 genes were downregulated in peri-infarct regions compared to equivalent regions of the contralateral hemisphere. These genes were primarily involved in mitochondrial respiration and neuronal function. Specifically, genes involved in mitochondrial fission and respiration, including *Cyc1*, *Coa3*, *Cox5a*, and *Mff*, and the neurofilament subunits *Nefh*, *Nefm*, and *Nefl*, were downregulated (**Fig. 10D**). Together, these results demonstrate that the transcriptome of the peri-infarct region is characterized by the upregulation of genes involved in inflammation and lipid metabolism and the downregulation of genes involved in mitochondrial respiration and neuronal function.

Next, we performed genome-wide expression analysis of the RNA-Seq data using GSEA. GSEA revealed significant enrichment of various biological processes in peri-infarct regions of aged mice, including (i) macrophage activation, (ii) cytokine production, (iii) BCR signaling pathway, (iv) lipoprotein metabolism, and (v) phagocytosis. In contrast, GSEA identified biological processes more significantly enriched in equivalent regions of the contralateral hemisphere, including (i) synaptic vesicle localization, (ii) mitochondrion organization, (iii) neurotransmitter secretion, (iv) myelin sheath, and (v) mitochondrial transport (**Fig. 11**). These results further indicate that peri-infarct regions are defined by chronic inflammation and dysregulated lipid metabolism, whereas equivalent regions of the contralateral hemisphere are defined by intact neuronal function.

**Figure 11.**
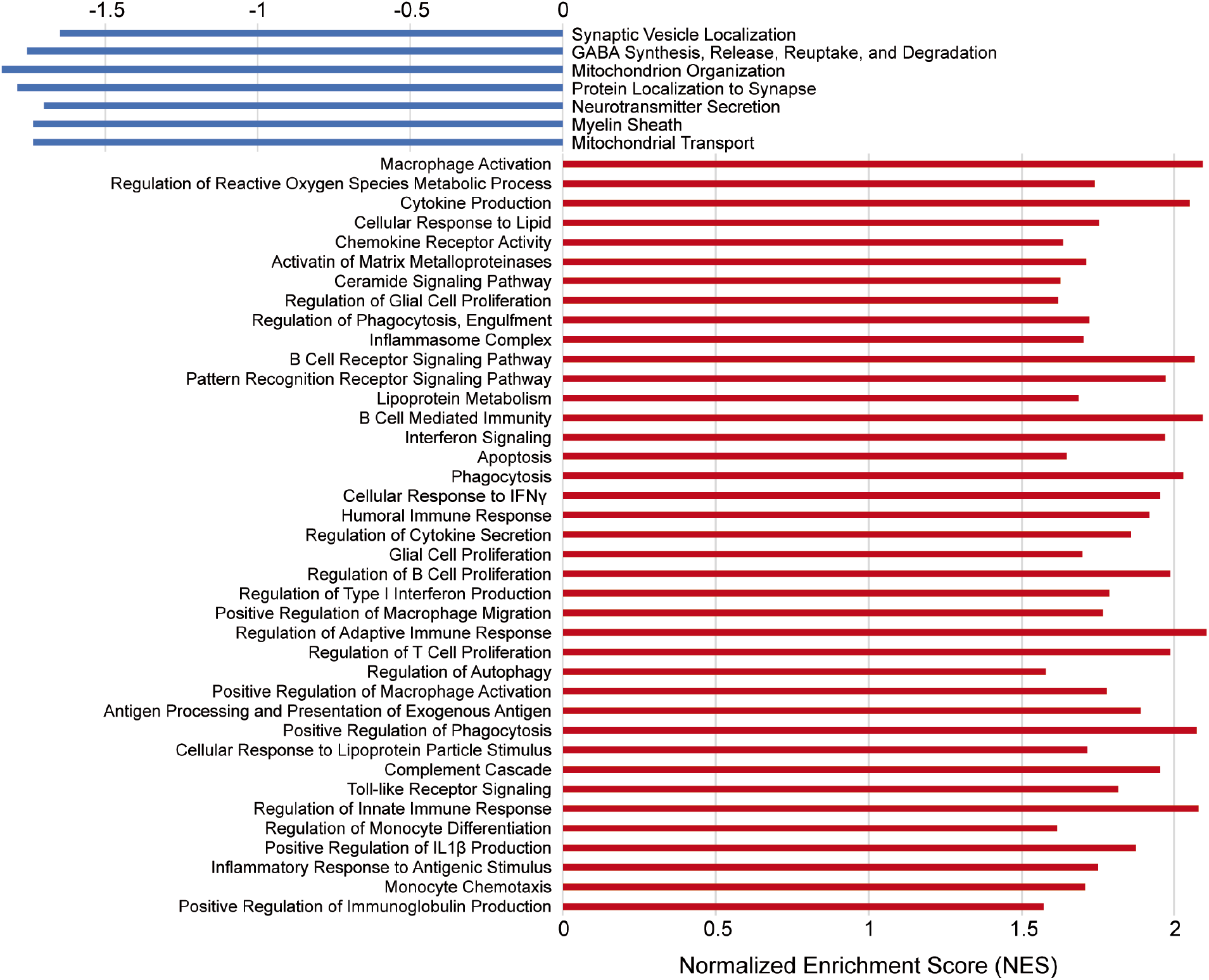
Biological processes enriched in peri-infarct regions of aged mice at 7 weeks after stroke. Gene set enrichment analysis (GSEA) revealed significant enrichment of multiple biological processes in peri-infarct regions of aged mice, depicted in red. Conversely, GSEA identified biological processes more significantly enriched in equivalent regions of the contralateral hemisphere, depicted in blue.

### HPβCD attenuates inflammation and increases transcripts associated with neuronal function in peri-infarct regions of aged mice after stroke

To evaluate the impact of HPβCD treatment on the transcriptome of the peri-infarct region, we performed bulk RNA-Seq on brain tissue collected 7 weeks after stroke. For this analysis, we compared the peri-infarct regions of vehicle- and HPβCD-treated aged mice (**Fig. 12A**). The peri-infarct region of HPβCD-treated mice was characterized by 5,385 differentially expressed genes (**Fig. 12B, C**). The DE analysis revealed that specific markers of reactive astrogliosis, including *Gfap*, *Serpina3n*, and *Stat3*, were downregulated following HPβCD treatment. The peri-infarct region of HPβCD-treated mice was also characterized by the downregulation of multiple degradative enzymes, including *Mmp2*, *Mmp8*, and *Mmp14*; chemokine receptors, including *Ccr2*, *Cx3cr1*; and *Cxcr4*, and scavenger receptors, including *Cd36*, *Cd68*, and *Cxcl16* (**Fig. 12D**).

**Figure 12.**
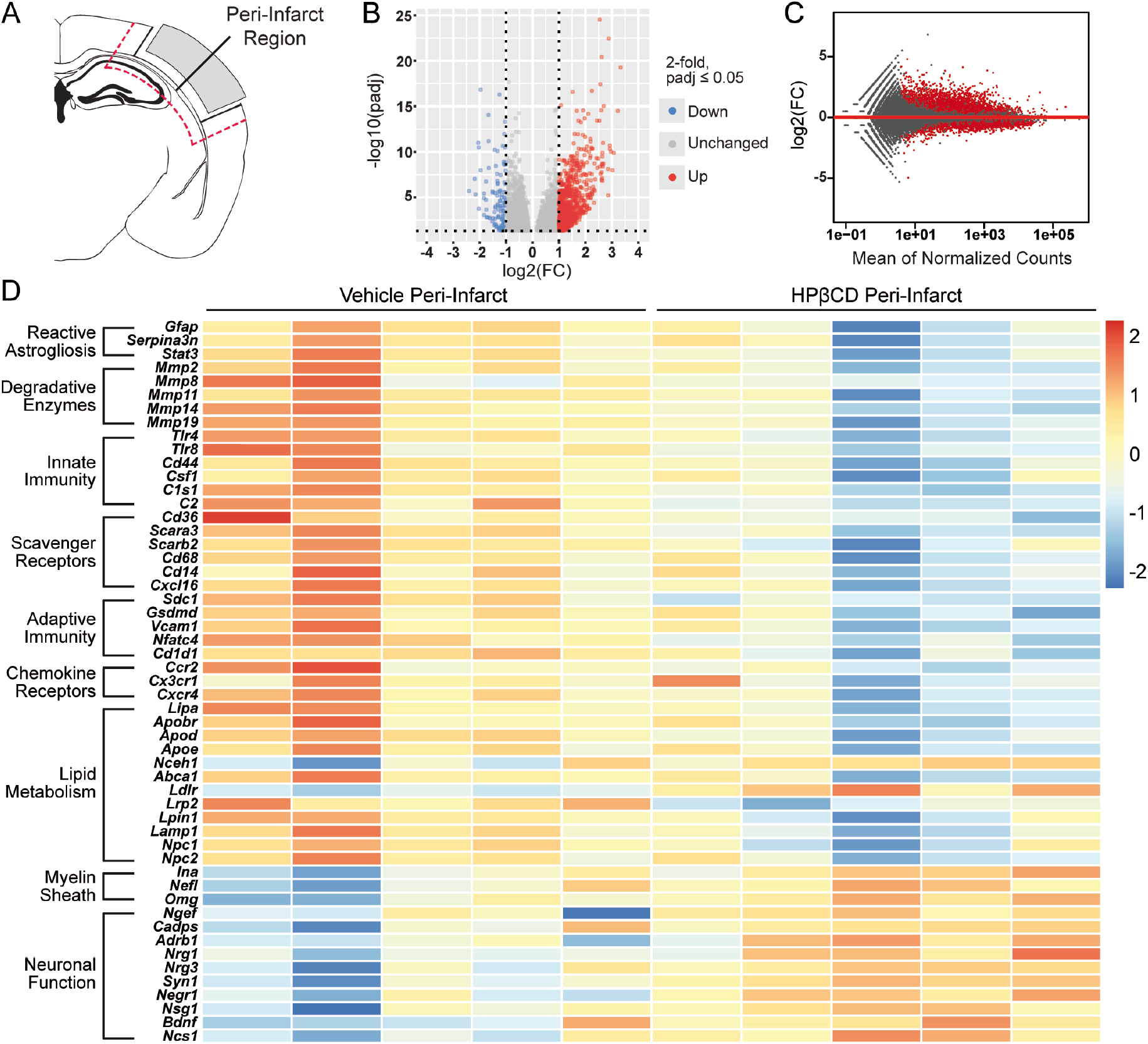
HPβCD attenuates inflammation and increases transcripts associated with neuronal function in peri-infarct regions of aged mice after stroke. ***A,*** Schematic of a mouse coronal brain section following stroke induced by distal middle cerebral artery occlusion + hypoxia, with the analyzed region indicated. ***B, C,*** Volcano (***B***) and MA (***C***) plots constructed from count data show differences in gene expression between peri-infarct regions from vehicle- and HPβCD-treated aged mice at 7 weeks after stroke (false discovery rate-adjusted p < 0.05; FC > |2|). ***D,*** Row-scaled heatmap displaying differentially expressed genes associated with lipid metabolism, inflammation, and neuronal function (false discovery rate-adjusted p < 0.05; FC > |2|). FC, fold change.

In addition, the transcriptomic analysis demonstrated a restoration of lipid homeostasis in peri-infarct regions of HPβCD-treated mice compared to vehicle-injected mice, as evidenced by the downregulation of *Lipa*, *Apoe*, *Lamp1*, *Npc1*, and *Npc2* and upregulation of *Nceh1* and *Ldlr*. A reduction in the innate immune response (identified by *Tlr4*, *Csf1*, and *C2*) and in the adaptive immune response (identified by *Sdc1*, *Gsdmd*, and *Cd1d1*) characterized peri-infarct regions of HPβCD-treated mice (**Fig. 12D**); these alterations are consistent with the attenuated chronic inflammation that characterized infarcts of the HPβCD-treated aged mice (**Fig. 8D**). These results demonstrate that HPβCD treatment after stroke attenuates chronic inflammation and promotes lipid metabolism in peri-infarct regions of aged mice. Additionally, the DE analysis revealed an upregulation of genes involved in myelin sheath structural integrity, such as *Ina*, *Omg*, and *Nefl*, and synaptic plasticity and neuroplasticity, such as *Syn1*, *Bdnf*, *Nrg1*, *Negr1*, and *Nsg1* (**Fig. 12D**). These results suggest that HPβCD treatment improves the structure and function of neurons in peri-infarct regions of aged mice after stroke.

Next, we performed genome-wide expression analysis of the RNA-Seq data using GSEA, which revealed a significant enrichment of various biological processes in peri-infarct regions of HPβCD-treated aged mice, including (i) synaptic vesicle localization, (ii) neuron projection membrane, (iii) dendritic spine organization, (iv) myelin sheath, and (v) regulation of axonogenesis. Conversely, biological processes more significantly enriched in vehicle-injected mice than in HPβCD-treated mice included (i) macrophage activation, (ii) binding and uptake of ligands by scavenger receptors, (iii) cytokine production, (iv) lipid droplet, (v) lymphocyte mediated immunity, and (vi) lipid catabolic process (**Fig. 13**). These results demonstrate that peri-infarct regions of HPβCD-treated aged mice are characterized by enhanced membrane integrity and reduced inflammation compared to vehicle-injected aged mice.

**Figure 13.**
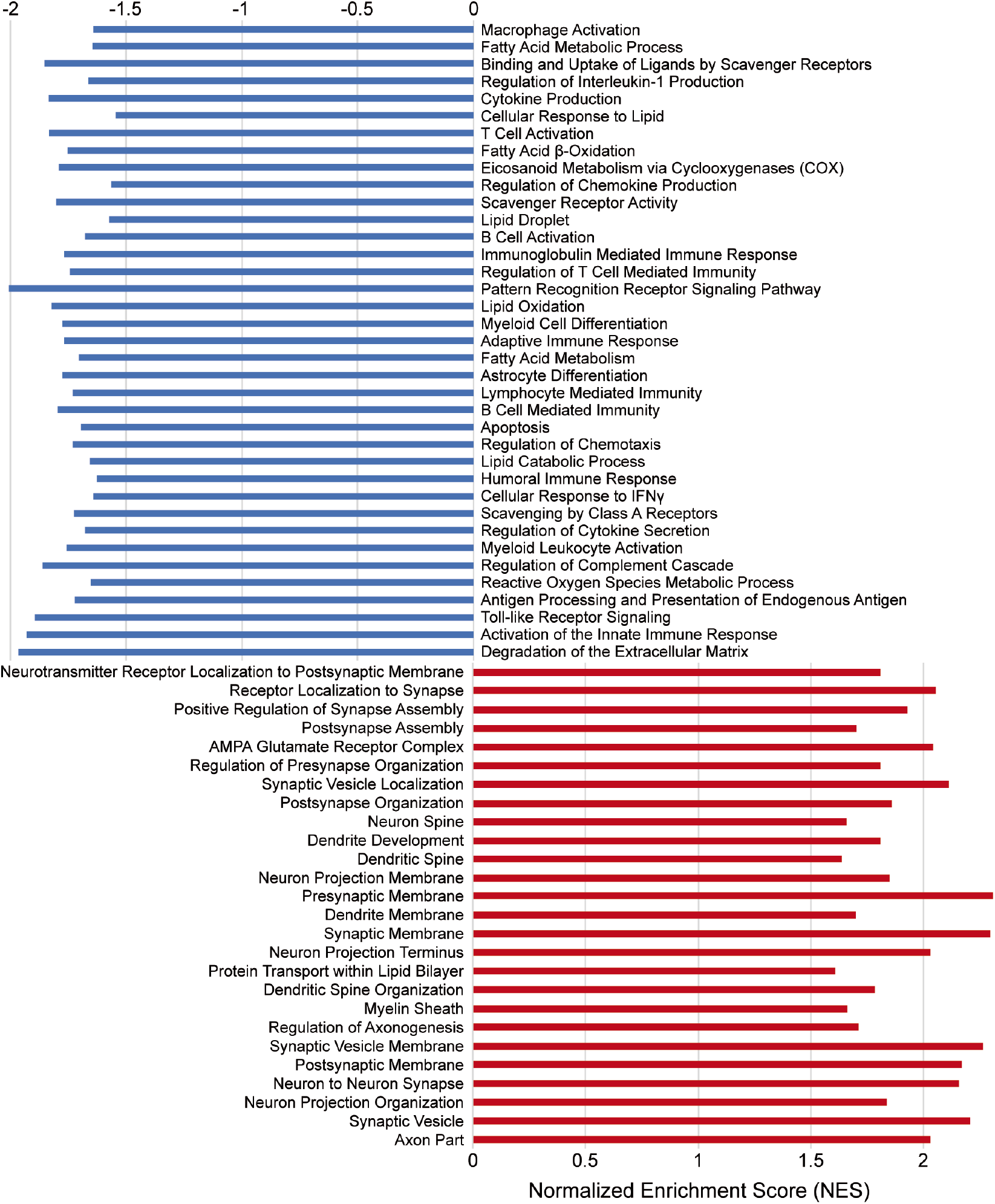
Biological processes enriched in peri-infarct regions of HPβCD-treated aged mice at 7 weeks after stroke. Gene set enrichment analysis (GSEA) revealed significant enrichment of multiple biological processes in peri-infarct regions of HPβCD- treated aged mice, depicted in red. Conversely, GSEA identified biological processes more significantly enriched in peri-infarct regions of vehicle-injected aged mice, depicted in blue.

### HPβCD attenuates neurodegeneration and improves recovery in aged mice after stroke

To assess behavioral outcomes following HPβCD treatment, we used the light/dark transition test and the Y-maze spontaneous alternation behavior (SAB) test. The light/dark transition test is based on the innate aversion of mice to brightly illuminated areas and on their spontaneous exploratory behavior in response to mild stressors (Bourin & Hascoët, 2003; Takao & Miyakawa, 2006). Our results revealed that in the light/dark box, HPβCD-treated mice spent less time in the brightly illuminated compartment than vehicle-injected mice (**Fig. 14A**). Additionally, HPβCD-treated mice were slower to emerge from the dark compartment after being habituated to the dark for 30 min prior to the test (**Fig. 14A**). These behaviors indicate decreased impulsivity and behavioral disinhibition in HPβCD-treated mice.

**Figure 14.**
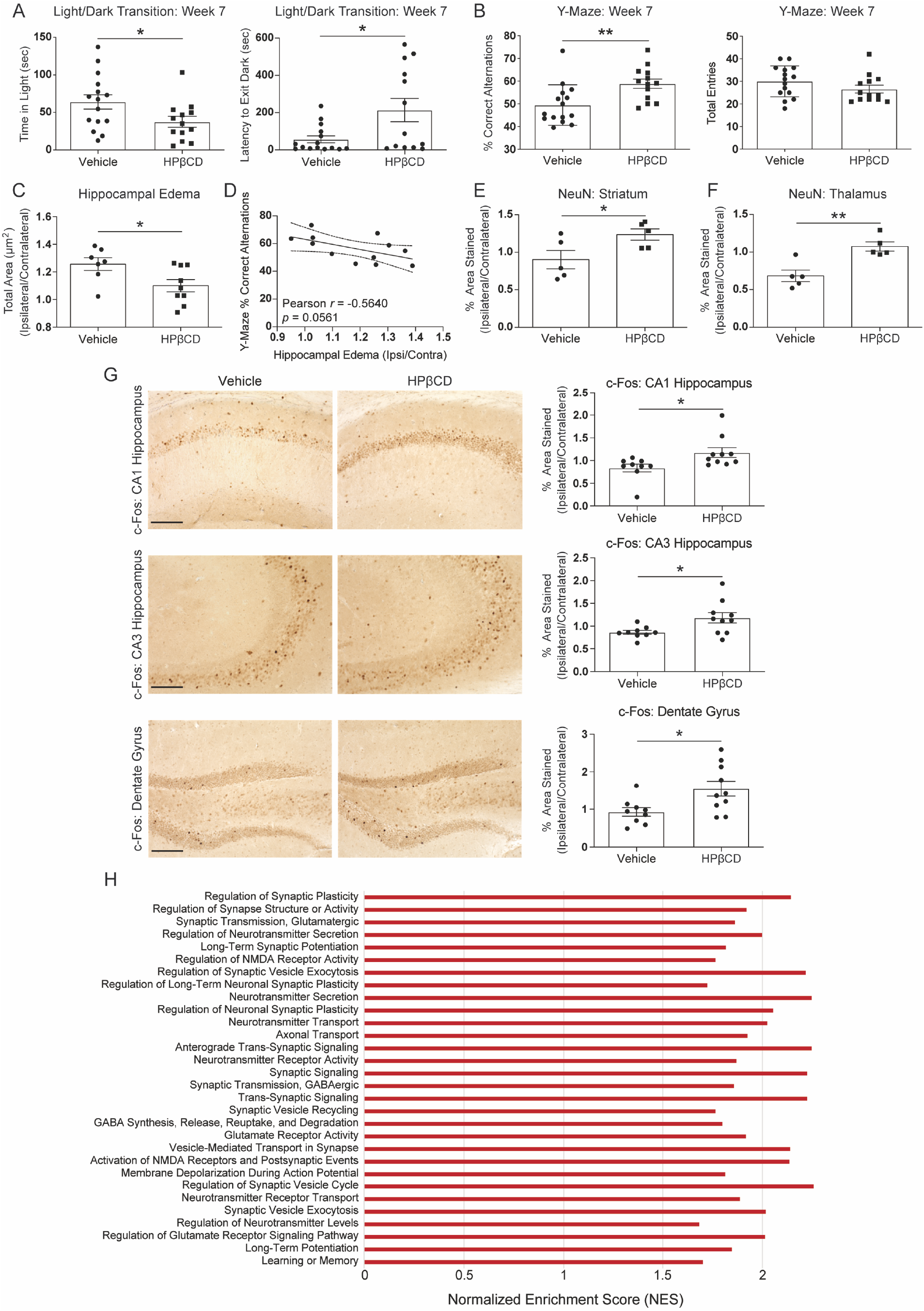
HPβCD attenuates neurodegeneration and improves recovery in aged mice after stroke. ***A,*** HPβCD-treated aged mice exhibited less impulsive behavior than vehicle-injected mice in the light/dark transition test at 7 weeks after stroke. (*n* = 13–15; unpaired *t* tests; \**p* < 0.05). ***B,*** HPβCD-treated aged mice exhibited intact spatial working memory in the Y-maze spontaneous alternation behavior test compared to vehicle-injected aged mice at 7 weeks after stroke. (*n* = 13–15; unpaired *t* tests; \**p* < 0.05). ***C,*** Quantification of hippocampal edema at 7 weeks after stroke in vehicle- and HPβCD-treated mice. (*n* = 7–9; unpaired *t* test; \**p* < 0.05). ***D,*** Pearson correlation between Y-maze spontaneous alternation behavior and hippocampal edema at 7 weeks post-stroke (*n* = 12; r = -0.5640; *p* > 0.05). ***E, F,*** Quantification of NeuN immunoreactivity in the striatum and thalamus from vehicle- or HPβCD-treated mice. (*n* = 5; unpaired *t* tests; \**p* < 0.05, \*\**p* < 0.01). ***G,*** Representative 20× images of c-Fos immunoreactivity in hippocampal regions from vehicle- or HPβCD-treated mice. Quantification of images is shown to the right of each photomicrograph. (*n* = 9–10; unpaired *t* tests; \**p* < 0.05, \*\*\**p* < 0.001). Scale bar, 125 μm. Data are presented as mean ± SEM. ***H,*** Gene set enrichment analysis revealed significant enrichment of biological processes associated with neuronal function and activity in peri-infarct regions of HPβCD-treated aged mice at 7 weeks after stroke.

We also assessed hippocampal-dependent spatial working memory at 7 weeks after stroke with the Y-maze SAB test (Yamada et al., 1996). We discovered that vehicle-injected mice developed a delayed cognitive deficit, whereas HPβCD-treated mice displayed neurotypical cognitive function comparable to baseline recordings (**Fig. 14B**). To assess hippocampal edema, we measured the total hippocampal area of the ipsilateral and contralateral hemispheres in aged mice that received vehicle or HPβCD. Hippocampal enlargement 7 weeks after stroke was attenuated in mice that had received HPβCD (**Fig. 14C**). Through Pearson’s correlation analysis, we observed a moderate negative correlation (*r* = -0.5640) between hippocampal edema and hippocampal-dependent spatial working memory outcomes in the Y-maze SAB test (**Fig. 14D**). We next evaluated secondary neurodegeneration by quantifying NeuN immunoreactivity as a biomarker of secondary neurodegeneration in regions of axonal degeneration. Repeated administration of HPβCD led to the preservation of NeuN immunoreactivity in the striatum and thalamus (**Fig. 14E, F**). These results demonstrate that HPβCD attenuates secondary neurodegeneration and improves cognitive function following stroke.

To investigate neuronal activation in hippocampal regions, we next measured the immunoreactivity of c-Fos, a transcription factor thought to primarily reflect NMDA- mediated neuronal activity (Albertini et al., 2018). We observed an induction of c-Fos expression in the dentate gyrus, CA1, and CA3 areas of HPβCD-treated mice (**Fig. 14G**). Importantly, Fleischmann et al. demonstrated a critical role for c-Fos in hippocampal-dependent spatial learning and memory, as well as in NMDA receptor-dependent LTP formation (Fleischmann et al., 2003). Together, our results show that an improvement in hippocampal-dependent spatial working memory (**Fig. 14B**) is associated with an induction of c-Fos in hippocampal regions of HPβCD-treated mice (**Fig. 14G**) at 7 weeks after stroke.

To corroborate these findings, we further analyzed the results from the GSEA performed on peri-infarct regions from vehicle- and HPβCD-treated aged mice (**Fig. 13**). We discovered that peri-infarct regions of HPβCD-treated aged mice were characterized by enriched biological processes that signify improved neuronal and synaptic function, including (i) long-term synaptic potentiation, (ii) neurotransmitter transport, (iii) synaptic signaling, (iv) glutamate receptor activity, (v) membrane depolarization during action potential, and (vi) learning or memory (**Fig. 14H**). These results provide evidence that HPβCD improves neuronal integrity in peri-infarct regions by 7 weeks after stroke.

## Discussion

Dysregulated lipid homeostasis has been implicated in chronic neurodegenerative diseases, including Alzheimer’s disease, Parkinson’s disease, and Niemann–Pick disease type C, and in acute neuronal injuries, including ischemic stroke (Vance, 2012). Lipid metabolism is particularly important in the mammalian brain, as lipids constitute 50–60% of the dry weight (Luchtman & Song, 2013). Notably, the mammalian brain is highly enriched in cholesterol, which primarily exists in two distinct pools within the CNS: in plasma membranes of glial cells and neurons and in myelin sheaths surrounding axons. Due to the intact blood–brain barrier, cholesterol synthesis and lipoprotein transport are regulated independently of the peripheral circulation (Dietschy & Turley, 2001; Vance, 2012). Therefore, further investigation into the role of altered lipid metabolism in neurodegeneration will likely aid in the development of novel therapeutic strategies.

Our previous preclinical studies indicate that there is substantial overlap between the pathological characteristics of chronic stroke infarcts and the hallmark features of atherosclerotic plaques. Similar to atherosclerotic plaques, chronic stroke infarcts contain foamy macrophages, lipid droplets, and intracellular and extracellular cholesterol crystals (Chung et al., 2018). We have also shown that the development of these pathologies coincides with the recruitment and infiltration of adaptive immune cells (Doyle et al., 2015; Zbesko et al., 2021). Importantly, in atherogenesis, the formation of foamy macrophages, caused by dysregulated lipid metabolic processes, leads to the recruitment of adaptive immune cells and the production of cytokines and degradative enzymes. Herein, we postulated that following stroke, lipids derived from myelin debris and other cell membranes overwhelm the processing capability of phagocytes in the brain, leading to the formation of lipid-laden foam cells, generation of cholesterol crystals, secretion of pro-inflammatory cytokines, and production of degradative enzymes. Further, we hypothesized that this chronic inflammatory response, coupled with concurrent cell death, causes post-stroke secondary neurodegeneration and impairs functional recovery.

Using the distal middle cerebral artery occlusion + hypoxia mouse model of stroke, we first characterized the lipidome of chronic stroke infarcts. The lipidome of infarcts at 7 weeks after stroke showed a substantial elevation of lipids, including sphingomyelins, sulfatides, and cholesterol esters, compared to contralateral cortices. We also observed an accumulation of intracellular lipid droplets. It has been shown that the excessive accumulation of cholesterol esters in lipid droplets contributes to foam cell formation (Yu et al., 2013). These findings suggest that the foam cells identified in chronic stroke infarcts contain an abundance of cholesterol and other myelin-derived lipids.

Using transcriptomic analyses, we assessed the impact of lipid dyshomeostasis on chronic stroke infarcts and peri-infarct regions. We anticipated disparity between ipsilateral and contralateral regions. Indeed, the transcriptomes of chronic stroke infarcts and peri-infarct regions were characterized by an upregulation of genes involved in inflammation, reactive astrogliosis, and lipid metabolism. Specifically, the upregulation of TLRs, including *Tlr1*, *Tlr2*, and *Tlr4*; scavenger receptors, including *Scarb2*, *Cd68*, *Msr1*, and *Cxcl16*; and other lipid mediators, including *Npc1*, *Npc2*, *Lamp1*, *Ffar1*, *Pparg*, *Apoe*, *Trem2*, and *Abca1* indicates a pronounced disruption in lipid homeostasis at 7 weeks after stroke. These transcriptomic analyses suggest that the mechanism of foam cell formation in ischemic stroke shares commonalities with the mechanism extensively characterized in atherosclerosis, albeit distinct in that myelin debris, rather than circulating LDL cholesterol, appears to be the primary source of lipids that overwhelm the processing capacity of infiltrating macrophages and resident microglia.

We next assessed the efficacy of HPβCD in counteracting or reversing foam cell formation and immune cell infiltration following stroke, as HPβCD induces liver X receptor target gene expression in macrophages and leads to an increase in cholesterol transporters that further promote cholesterol efflux (Zimmer et al., 2016). Cyclodextrins are cyclic oligosaccharides comprised of glucose monomers. With hydrophobic interiors and hydrophilic exteriors, cyclodextrins can form inclusion complexes with hydrophobic compounds and yield aqueous solubility and stability across tissues. Cyclodextrins are commonly utilized as carriers and solubilizing agents for steroids, antivirals, and chemotherapies (Gidwani & Vyas, 2015; Rasheed et al., 2008). In addition, HPβCD has also proven efficacious in the regression of atherosclerotic plaques and the prevention of age-related lipofuscin accumulation (Gaspar et al., 2017; Zimmer et al., 2016). Here, we found that repeated administration of HPβCD after stroke significantly attenuated chronic inflammation in young adult and aged mouse models. Specifically, there were substantially fewer intracellular lipid droplets, B lymphocytes, T lymphocytes, and antibody-producing plasma cells in the infarcts of young adult and aged HPβCD-treated mice than in vehicle-injected mice at 7 weeks after stroke.

The discovery that HPβCD is effective in attenuating B-lymphocyte accumulation in chronic stroke infarcts is significant because we have shown that B lymphocytes mediate cognitive dysfunction following stroke (Doyle et al., 2015). Correspondingly, activated B lymphocytes and autoantibodies have been shown to cause neuropathology in models of experimental autoimmune encephalomyelitis (EAE), SCI, and transient middle cerebral artery occlusion (Ankeny et al., 2006; Ortega et al., 2015; Raine et al., 1999). However, the B-lymphocyte response to stroke is heterogeneous and has also been shown to be beneficial for stroke recovery. For example, it consists of both T-lymphocyte dependent and T-lymphocyte independent responses (Weitbrecht et al., 2021; Zbesko et al., 2021). Additionally, B-lymphocyte depletion following stroke reduces stroke-induced hippocampal neurogenesis and impairs functional recovery (Ortega et al., 2020). Future studies are necessary to elucidate the role of infiltrating B lymphocytes in stroke recovery.

Using transcriptomic analyses, we next assessed transcripts in infarct and peri-infarct regions of HPβCD-treated aged mice. We found that systemic administration of HPβCD distinctively altered the transcriptomes of infarct and peri-infarct regions. In infarcts, HPβCD upregulated genes involved in lipid metabolic processes, including *Lipa*, *Npc1*, *Apoe*, and *Abca1*; in peri-infarct regions, HPβCD had an inverse effect on these genes. These observations suggest that HPβCD has supported the restoration of lipid homeostasis through transcriptional reprogramming of macrophages, consistent with a previously described mechanism of action (Zimmer et al., 2016). In addition, HPβCD influenced expression profiles in infarcts through the upregulation of genes involved in angiogenesis and in peri-infarct regions through the upregulation of genes involved in neuronal and synaptic structure and function. Together, these results indicate that HPβCD exerts multiple restorative effects on affected brain regions after stroke.

We next evaluated the efficacy of HPβCD in enhancing recovery after stroke. We assessed impulsivity and risk-taking behavior using the light/dark transition test. We have previously demonstrated a chronic impact of stroke on the psychological measurement of impulsivity in the light/dark transition test, resembling the uninhibited, risk-taking behaviors exhibited by patients with Alzheimer and frontotemporal dementias (Liscic et al., 2007; Nguyen et al., 2018). HPβCD-treated mice spent less time in the brightly illuminated chamber and were slower to exit the dark chamber. These behaviors suggest that HPβCD-treated mice act less erratically and impulsively than vehicle-injected mice. We also measured hippocampal-dependent spatial working memory using the Y-maze spontaneous alternation behavior test. We found that vehicle-injected mice developed a delayed cognitive deficit, whereas HPβCD-treated mice displayed neurotypical cognitive function. This improvement in recovery was associated with a reduction in hippocampal edema, a preservation of neurons in the striatum and thalamus, and an induction of c-Fos expression in hippocampal regions (dentate gyrus, CA1, CA3). Correspondingly, peri-infarct regions of HPβCD-treated mice were defined by an enrichment of biological processes signifying improved neuronal and synaptic integrity. Thus, administration of HPβCD reduced stroke-induced neuropathology and improved cognitive function after stroke.

For decades, putative neuroprotectants have failed in clinical stroke trials despite demonstrating efficacy in preclinical studies. In light of these failures, we followed Stroke Therapy Academic Industry Roundtable (STAIR) guidelines to accelerate the translation of HPβCD from bench to bedside (Landis et al., 2012; Lapchak et al., 2012; STAIR, 1999): (i) we performed and analyzed experiments in a randomized and blinded manner; (ii) we calculated sample sizes based on a power analysis prior to the start of the study; (iii) we delayed treatment by 1 week to achieve an optimal therapeutic time window; (iv) we selected a dosage and a systemic route of administration to parallel preclinical trials using HPβCD in Niemann–Pick disease type C; (v) we chose a model of stroke in which long-term functional outcomes could be assessed; and (vi) we utilized both young adult and aged animals. However, there are additional experiments that need to be conducted. For example, we have not yet tested the efficacy of HPβCD in female mice. Although we anticipate comparable efficacy in females, future studies are needed to address sex as a biological variable. In addition, HPβCD must still be tested in a model of transient occlusion, in a different laboratory, and in a second species.

In conclusion, we have shown that, coincident with the progressive infiltration of adaptive immune cells, chronic stroke infarcts accumulate lipids, including sphingomyelins, sulfatides, and cholesterol esters. To our knowledge, this substantial disruption in lipid homeostasis has not been previously recognized or investigated in the context of stroke. We also discovered that repeated administration of HPβCD following stroke aids in the restoration of lipid homeostasis, attenuates lipid droplet and immune cell accumulation, and improves recovery at transcriptional and functional levels. Therefore, HPβCD warrants further testing to determine if it can be repurposed for the treatment of ischemic stroke and other CNS injuries.

## Acknowledgments

This work was funded by NINDS R01NS096091 (KPD), NIA R01AG063808 (TVN), NIA R21AG062781 (TVN), United States Department of Veterans Affairs I01RX003224 (RGS), NIA T32AG058503-01A1 (DAB), NINDS F31NS105455 (JCZ), and the Fondation Leducq Transatlantic Network of Excellence Stroke-IMPaCT (KPD). We thank Dr. Oswald Quehenberger (UCSD Lipidomics Core), Branden Lau, and Jonathan Galina-Mehlman (University of Arizona Genetics Core) for their technical assistance with lipidomics and transcriptomics analyses. We also thank Dr. Julia Slone-Murphy (NeuroEdit Ltd) for editing a draft of this manuscript.

